# New Vista into Origins of Viruses from a Prototypic ssDNA Phage

**DOI:** 10.1101/2022.05.27.493684

**Authors:** Nejc Kejzar, Elina Laanto, Ilona Rissanen, Vahid Abrishami, Muniyandi Selvaraj, Sylvain Moineau, Janne Ravantti, Lotta-Riina Sundberg, Juha T. Huiskonen

**Author notes:** Krupic lab, Department of Physiology, Development and Neuroscience, University of Cambridge, Cambridge CB2 3EG, UK. The Sainsbury Laboratory, Norwich Research Park, Norwich NR4 7UH, UK. contributed equally (Nejc Kejzar, Elina Laanto, Ilona Rissanen).

## Abstract

Viruses play a central role in all ecological niches; the origin of viruses, however, remains an open question. While phylogenetic analysis of distantly related viruses is hampered by a lack of detectable sequence similarity, structural biology can reveal conserved capsid protein structures that facilitate the study of distant evolutionary relationships. Here, we characterize the lipid-containing ssDNA temperate bacteriophage ΦCjT23, which is infecting *Flavobacterium* sp. (*Bacteroidetes*). We further detected ΦCjT23-like sequences in the genome of strains belonging to several *Flavobacterium* species. The virion structure determined by cryogenic electron microscopy revealed similarities to members of the viral kingdom *Bamfordvirae* that currently consists solely of dsDNA viruses. Common to these viruses, infecting hosts from all domains of life, is a major capsid protein composed of two upright β-sandwiches. The minimalistic structure of ΦCjT23 suggests that this phage serves as a model for the last common ancestor between ssDNA and dsDNA viruses in the *Bamfordvirae*. Both ΦCjT23 and the related phage FLiP infect *Flavobacterium* species found in several environments, suggesting that these types of viruses have a global distribution and shared evolutionary origin. Detailed comparisons to related, more complex viruses not only expand our knowledge about this group of viruses but also provide a rare glimpse into early virus evolution.

## Introduction

Several competing theories exist for the emergence of viruses^1^. In the absence of fossils, the study of virus evolution is limited to the present-day viruses, which can nevertheless still provide valuable clues. From our exploration of the virosphere some common principles are emerging, even amongst seemingly unrelated viruses. While high-throughput genome sequencing of environmental samples has shown that a significant fraction of newly obtained viral sequences has no homology to any known viral genome^2,3^, comparative analysis of virus structures allows structure-based phylogenetics. With this approach, several viruses have been grouped into a limited number of viral lineages, where viruses belonging to a certain lineage often infect hosts from entirely different domains of life^4^.

One viral lineage that has emerged as a result of comparative structural virology studies is the PRD1–adenovirus lineage, consisting of several dsDNA viruses infecting hosts ranging from humans^5,6^ to eukaryotic unicellular algae^7^ as well as bacteriophages (phages) infecting Gram-negative^8,9^ and Gram-positive^10^ bacteria. These dsDNA viruses share the same major capsid protein (MCP) fold and have recently been classified as members of the kingdom *Bamfordvirae*^*11*^. PM2 (*Pseudoalteromonas virus PM2, Corticoviridae family*), a dsDNA phage infecting strains of *Pseudoalteromonas*, is considered to be the prototypic member of this kingdom due to its minimalistic MCP fold^12^. Earlier, we have reported a phylogenetic connection between phage PM2 and the ssDNA phage FLiP (*Flavobacterium virus FLiP, Finnlakeviridae*), which infects flavobacteria^13^. At the genomic level, the 9,174-nt-long circular FLiP genome and the 10,079-bp-long circular PM2 genome share only 43% identity. Structurally, however, the two share the same capsid organization, characterized by the triangulation (*T*) number pseudo *T*=21 *dextro*. Furthermore, their MCPs share the same conserved double-jelly-roll β-sandwich/β-barrel fold, a hallmark of *Bamfordvirae*^12,13^. Importantly, these findings may challenge the accepted view of using the virus genome type as the main high-level classification criterion^13^.

Despite extensive efforts to characterize the structural landscape of the *Bamfordvirae* viral kingdom^6-10,14-20^, structurally similar viruses with an alternative genome type still comprise only one representative, phage FLiP^13^. To increase our confidence in linking the evolutionary origins of ssDNA viruses and dsDNA viruses, more comparative structural studies between viruses harboring different genome types are required. Such analyses are crucial not only to better understand the constraints posed by the structural requirements for virion assembly, but also to provide new insights into the early evolution of viruses.

In this study, we report a detailed characterization of ΦCjT23, an icosahedral temperate ssDNA phage with an inner lipid membrane, isolated earlier from a strain of *Flavobacterium* (previously *Cytophaga*)^21^. The determined sequence of its relatively short (7,642 nt) circular ssDNA genome showed little similarity to any known virus. The MCP sequence-based analysis, however, revealed an abundance of ΦCjT23-like prophages in the genomes of several bacterial strains belonging to various *Flavobacterium* species. ΦCjT23 virion structure was determined by cryogenic electron microscopy (cryo-EM) of purified viral particles and revealed a capsid organization with triangulation number pseudo *T*=21 *dextro*, similar to phages PM2 and FLiP^12,13^. The MCP has a minimalistic architecture in comparison to other homologous MCPs. The capsid vertex architecture was found to be unique and comprised of a flexible pentameric spike protein that includes a non-canonical penton base domain, differing from the specialized penton proteins of *Bamfordvirae* members. We conclude by deriving a possible scenario for the evolution of ΦCjT23-like ssDNA and dsDNA viruses of *Bamfordvirae* from a shared ancestor with a primordial capsid.

## Results

### Temperate phage ΦCjT23 has a circular ssDNA genome

Phage ΦCjT23 was investigated as part of our wider efforts to better understand the virosphere of lipid-containing bacteriophages. Phage ΦCjT23 was purified from a lysate (∼1.1x10^13^ PFU) using polyethylene glycol precipitation and a two-step gradient centrifugation resulting to ∼6.6x10^8^ PFU of purified virions (yield less than 0.01%). Genome composition of ΦCjT23 was identified to be ssDNA by degradation of its nucleic acids with DNase and S1 nuclease, while RNase was ineffective. Prior to sequencing, the complementary strand was synthesized with Klenow fragment. Sequencing revealed a circular genome of 7,642 nt (30.5% GC) with 15 predicted open reading frames (ORFs) (Fig. 1; Extended Data Figure 1). BlastN search returned only one significant result; a highly similar nucleotide sequence detected in *Flavobacterium* sp. KBS0721 (73% query coverage, 89% identity). BlastP and HHPred searches detected no conserved domains (Supplementary Data Table 1). ORF3 received several hits in the GenBank protein database to replication initiation proteins, including the replication initiation factor in *Cellulophaga* phages with ssDNA genomes (phi18:4, phi12a:1, phi12:2). Results from the HHPred search for the ORF05 indicated homology with the MCP of *Flavobacterium* phage FLiP (probability 95.5%, E-value 1.2). In addition, hits showed similarity to the *Pseudoalteromonas* phage PM2 minor membrane proteins P3 (ORF6 in ΦCjT23, probability 47.2%, E-value 28) and P6 (ORF11, probability 45.3%, E-value 37). While these hits were statistically insignificant, they hinted at homology between ΦCjT23, FLiP and PM2, which was confirmed by comparing the structures of the three phages (see below).

**Figure 1.**
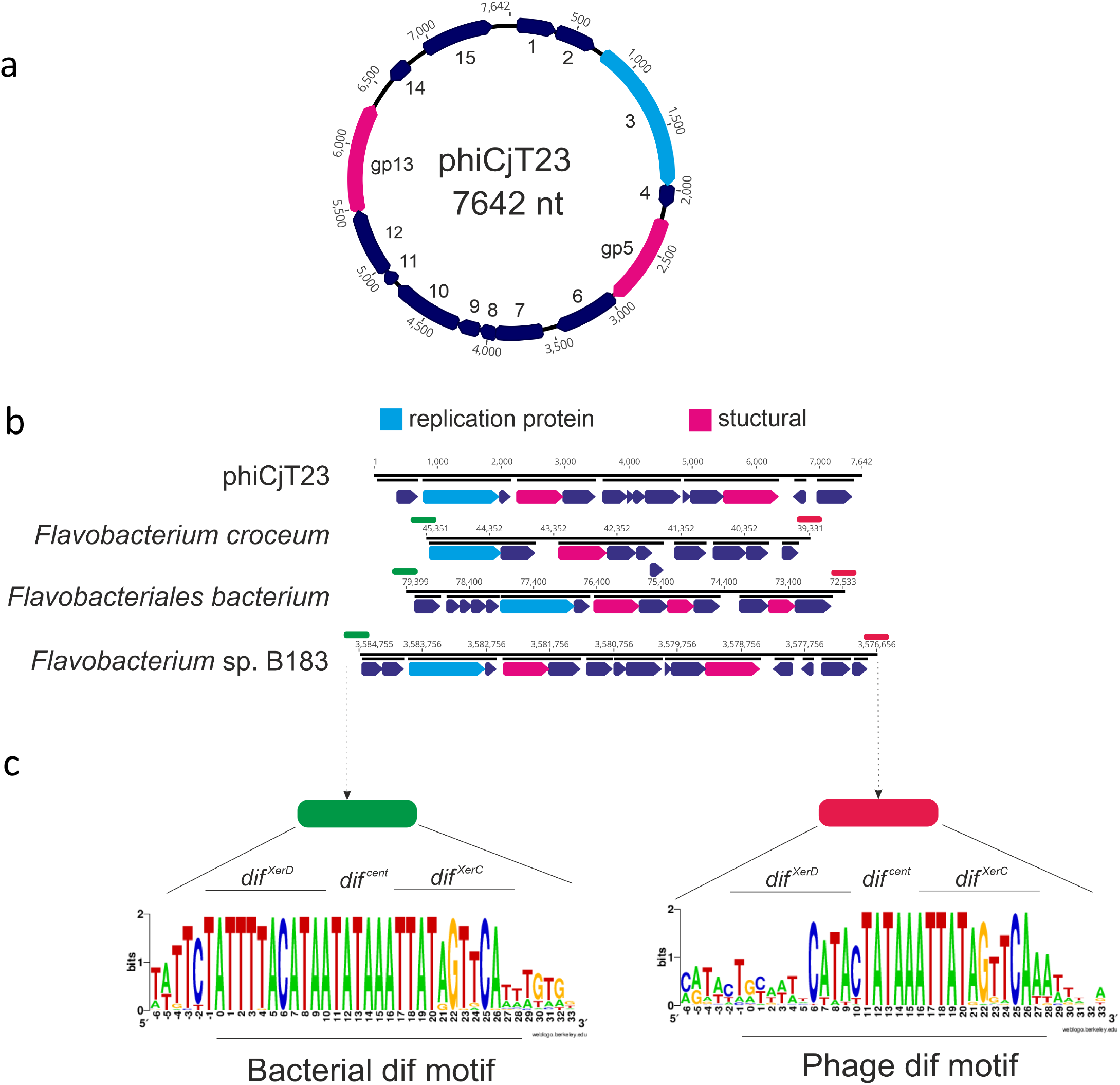
Genome of a temperate phage phiCjT23 and related prophages found in *Flavobacterium* genomes. **(a)** ΦCjT23 genome is a 7,642-nt long circular single-stranded DNA molecule with 15 predicted open reading frames (ORFs). The annotated replication initiation protein is indicated in blue and the two identified structural proteins, major capsid protein (MCP, gp5) and spike protein (gp13), are in pink. **(b)** Examples of putative prophages found in *Flavobacterium* genomes. The lengths of the sequences range between ∼6,000 and 8,000 bp. Shared genome synteny between the identified prophages included replication initiation protein near the start of the sequence, followed by a gene encoding a major capsid protein MCP (sequence similarity shared between the prophages). **(c)** Phages and bacterial dif-motifs were identified from the non-coding regions surrounding putative prophages implying XerC/XerD dependent integration. The 28-nt long dif locus (including six extra surrounding nucleotides) with XerC and XerD-binding sites, as well as the central hexanucleotide, are shown.

### ΦCjT23-like prophages detected in *Flavobacterium* genomes

BlastP searches of ΦCjT23 ORFs resulted in several hits against *Flavobacterium* genomes suggesting the presence of integrated ΦCjT23-like phages, or prophages, in bacterial genomes. In the set of 57 putative prophage sequences (ranging in size from 6,021 to 8,407 bp; Extended Data Figure 1; Supplementary Data Table 2) detected with this method, a gene coding for a replication initiator protein was followed in proximity by a gene coding for a putative MCP. BlastP search among the detected sequences yielded few hits to FLiP genome (Extended Data Figure 1). These hits were to gp16 which follows the predicted replication protein gp15 in FLiP genome. Similarly, as with ΦCjT23 genome, some of the detected sequences received hits to ssDNA phages infecting *Cellulophaga* (Extended Data Figure 1). Also, a putative *dif* site flanking both ends of these putative prophages was detected. A 28-nt long palindromic sequence was present both downstream and upstream of the prophage sequence, including motifs for *xerC* and *xerD* genes centered by a conserved hexanucleotide (Fig. 1, including also the surrounding six nucleotides). Several ssDNA phages, such as the filamentous Vibrio phage ΦCTX, use the *dif* site for integration into the host chromosome^22^. These results indicate that ΦCjT23-like phages use XerC/XerD mediated recombination for integration into *Flavobacterium* genomes.

### An icosahedral *T*=21 protein shell encloses a lipid membrane and the genome in the ΦCjT23 virion

To address distant evolutionary relationships, we determined the structure of the ΦCjT23 virion by cryo-EM (Fig. 2; Extended Data Figure 2). Spherical particles approximately 60 nm in diameter and homogenous in shape were observed in the micrographs (Extended Data Figure 2a). Occasionally, smaller particles were present, possibly corresponding to particles lacking the capsid and containing only the inner lipid envelope with the genome (Extended Data Figure 2a,b). Single-particle analysis of complete particles resulted in an icosahedrally symmetric reconstruction with the average estimated resolution of 4.1 Å. Spike proteins extend from the twelve vertices, but the density corresponding to these was disordered in the icosahedrally averaged map (Fig. 2a; Extended Data Figure 2c) due to the presence of a flexible hinge region in the spike. The basic capsid building block is the pseudo-hexameric MCP trimer (Fig. 2b). The observed 200 MCPs, occupying four unique positions in the asymmetric unit, follow icosahedral capsid organization characterized by a pseudo-triangulation number *T*=21. Within the asymmetric unit, type 1 trimers circle the five-fold symmetry axes at the vertices, type 2 trimers reside on both sides of the icosahedral two-fold axis of symmetry, type 3 trimers reside on the icosahedral three-fold axis of symmetry and type 4 trimers reside between the rest. This capsid organization is also found in dsDNA phage PM2 and ssDNA phage FLiP^12,13^. The dimensions of the faceted capsid are characterized by a face-to-face distance of ∼55 nm, an edge-to-edge distance of ∼54 nm, and a vertex-to-vertex distance of ∼63 nm. Inside, a lipid bilayer with a thickness of ∼5 nm is discernible (Fig. 2a; Extended Data Figure 2c). Weak density hints at minor structural components under trimers 2 and 4, possibly anchoring the lipid bilayer to the capsid (Fig. 2a).

**Figure 2.**
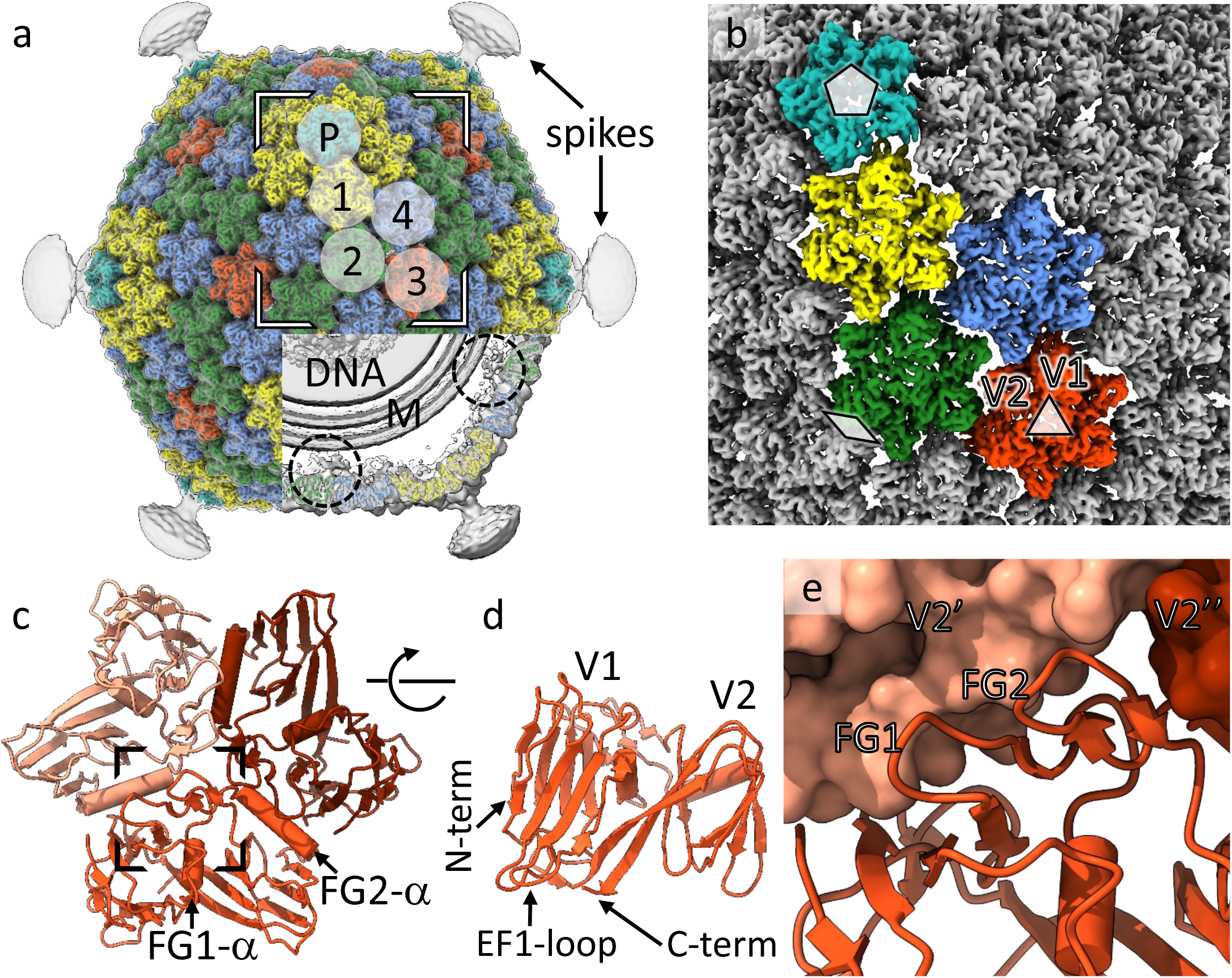
Structure of the ΦCjT23 bacteriophage capsid and major capsid protein trimer. **(a)** Surface rendering of the ΦCjT23 virion cryo-EM map at 4.1 Å resolution. The four different types of major capsid protein trimers are colored yellow (1), green (2), red (3) and blue (4). The penton base (P) is colored in cyan with the remaining structural components colored gray. On top, a gray transparent surface, calculated from the cryo-EM map filtered to 12-Å resolution, is shown. This lower-resolution surface reveals disordered density attributed to flexible spikes which originate from the penton bases. A part of the density is removed, to reveal the lipid bilayer (M), DNA and putative minor protein components (circles) bridging MCPs to the lipid bilayer. **(b)** A close-up of the high-resolution surface from the area indicated in *a* and rotated to face the viewer. The icosahedral axes of symmetry are marked with a pentagon (five-fold axis), triangle (three-fold axis) and diamond (two-fold axis). Type 1 trimers are peripentonal, circling the five-folds. Type 2 trimers are adjacent to the two-folds. Type 3 trimers locate on the three-folds. Type 4 trimers are farthest from any symmetry axes. **(c)** Model for one trimer (type 3) is shown as a ribbon. The three different subunits are shown in different shades. **(d)** The view in *c* is shown after rotating 90 degrees around the horizontal axis as indicated. Only the frontmost monomer is shown. The two β-sandwich domains (V1 and V2) are labeled, in addition to the protein termini. **(e)** Closeup of the area indicated in *c* show how the two upright loops (FG1 and FG2) lock the monomer with the V2 domains (V2’ and V2’’) of its neighbors (shown as surfaces).

### The ΦCjT23 major capsid protein is a canonical upright double-β sandwich

We determined the atomic model of ΦCjT23 MCP to compare with previously determined MCP structures. We used localized reconstruction^23^ to determine a density map for each trimer type (see Methods; Extended Data Figure 2d,e). This allowed refining local distortions in the capsid structure, improving the resolution of all trimers from 4.1 Å to 3.2 Å (FSC at 0.143 threshold; Extended Data Figure 3a). The improved map quality was sufficient to build atomic models of the trimers *de novo* (Extended Data Figure 3b–d). The entire polypeptide sequence corresponding to ORF5 (M1–V239) fitted the density, confirming this ORF as the MCP (P5).

**Figure 3.**
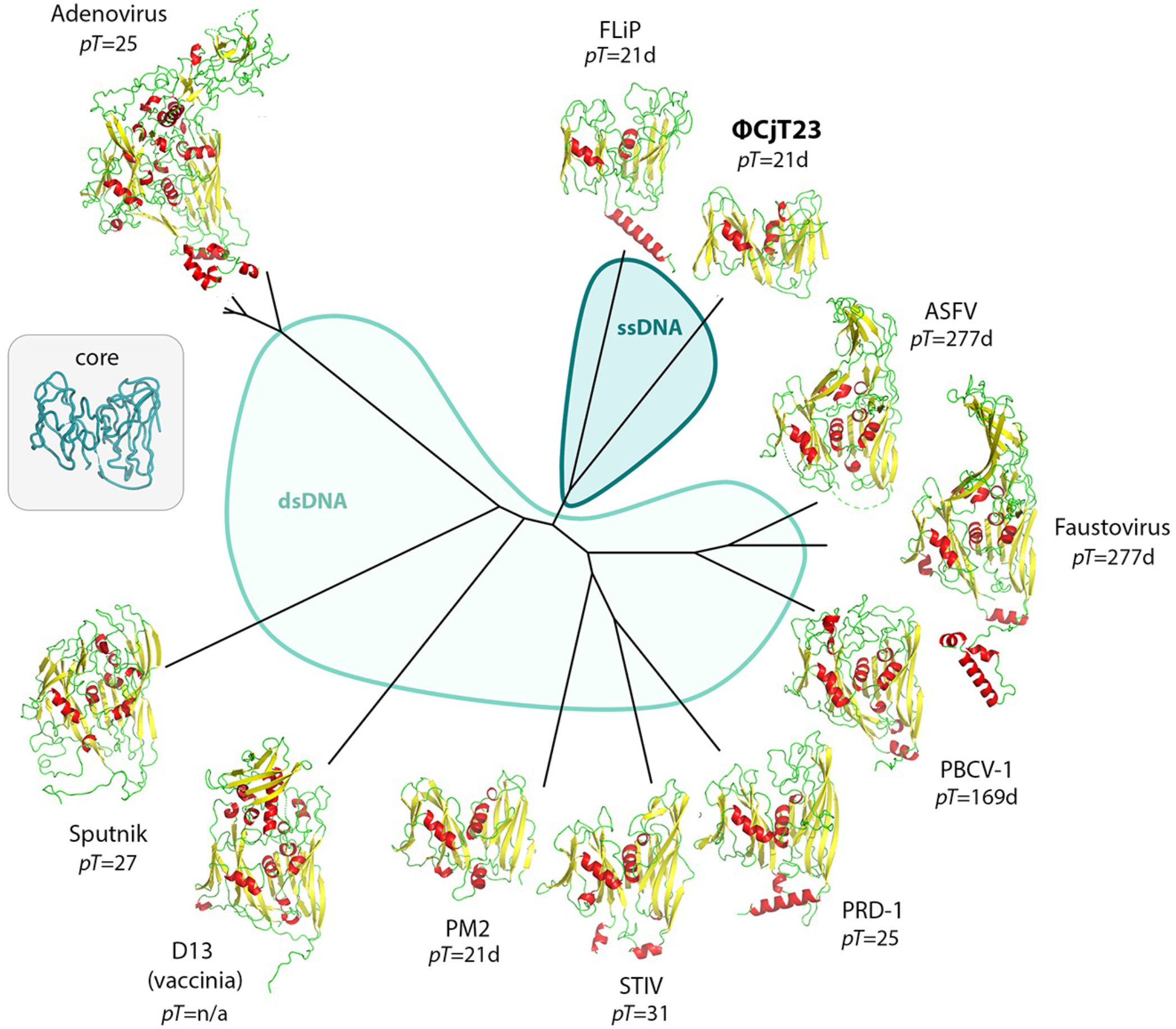
Structural phylogeny of major capsid proteins with double-jelly-roll fold. The two different parts of the phylogeny, corresponding to viruses harboring either an ssDNA (blue) or dsDNA (gray) genome, are shown in different colors. A generalized common core of the fold used in the comparisons is shown as a C_α_-trace in the middle. The *T*-number for the corresponding icosahedral capsid, when applicable, is given.

The MCP monomer is composed of two β-sandwiches, V1 (M1–Q126) and V2 (T127– V239). Both domains follow a classic upright jelly-roll topology^5^, each of which is composed of two antiparallel β-sheets composed of four strands, denoted B-I-D-G and C-H-E-F, respectively^24^ (Fig. 2d; Extended Data Figure 4). The β-strands are connected by loops. The BC, DE, FG, and HI loops are facing the capsid exterior, whereas CD, EF, and GH loops, in addition to N- and C-termini, are facing the capsid interior. Unique to ΦCjT23, the FG loops (FG1 and FG2) stand upright and seem to stabilize the trimer by interacting across subunit boundaries (Fig. 2e). ΦCjT23 MCP also contains two α-helices, a characteristic to all MCPs of the *Bamfordvirae*. These helices are embedded in the FG loop region in both V1 and V2 domains (termed FG1-α in V1 and FG2-α in V2). FG1-α is lodged between V1 and V2 domains of each monomer, whereas FG2-α resides between V2 and V1 of the adjacent monomer of the same trimer (Fig. 2c).

**Figure 4.**
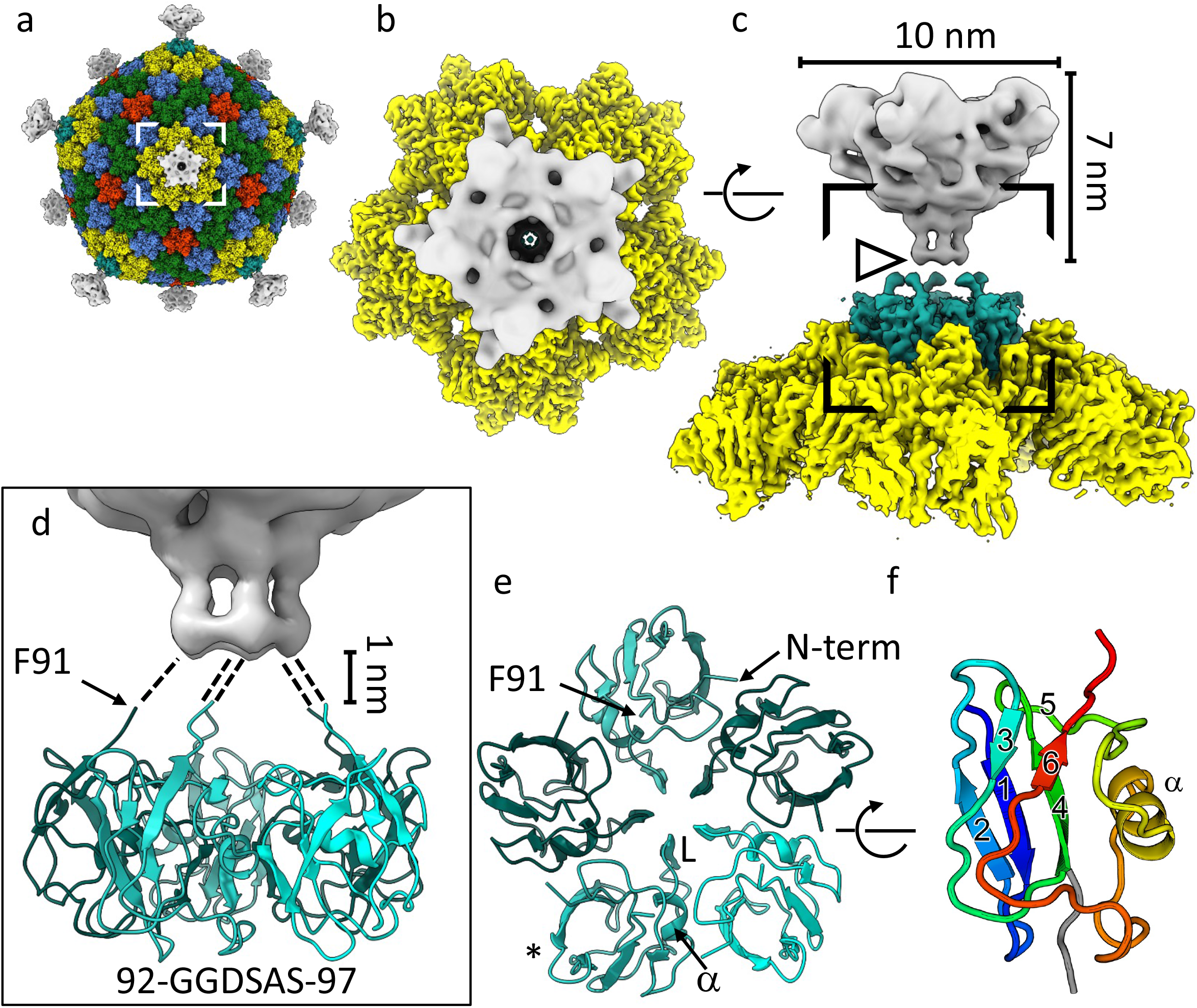
ΦCjT23 vertex. **(a)** A composite cryo-EM map of ΦCjT23 bacteriophage capsid combining localized reconstructions of the four MCP trimers (yellow, green, red and blue), penton base (cyan) and spike (gray). **(b)** A close-up of one vertex seen from the top from the area indicated in *a*. (**c**) The same view as in *b*, rotated by 90 degrees around the axis indicated. The dimensions of the spike, corresponding to the external domains of spike protein P13, are indicated. Note the discontinuation in the density (arrowhead). **(d)** A close-up of the area indicated in *c*. The model of the penton base part of the spike is shown in ribbon (cyan). The ordered part of the map ends around the area occupied by G92. Putative linker regions spanning the 1-nm gap between the penton base and external part of the spike (gray) are illustrated with dashed lines. The sequence of the putative linker is given. **(e)** A model of the pentamer formed by the penton base domain seen along a 5-fold axis of symmetry. Different monomers are colored with different shades of cyan. The positions of the N-terminus, C-terminal G92, αhelix at the monomer– monomer interface (α) and loop (L) closing the oculus in the middle of the pentamer are labelled. **(f)** The monomer is colored from blue (N-terminus) to red (C-terminal G92) marked with an asterisk in *e* is shown rotated by 90-degrees around the axis indicated. The β-strands are numbered 1–6. Strands 2-1-4 make a β-sheet. The α-helix, residing in the region connecting strands 5 and 6, is labeled.

### A simple molecular switch at the MCP trimer–trimer interface

How does the pseudo *T*=21 capsid assemble? The asymmetric unit contains ten independent MCP monomers, contributing a total of 20 β-sandwiches (V1 and V2). The penton domain of the spike presents one monomer per asymmetric unit and constitutes an additional building block (Fig. 2b). As these 21 building blocks are not identical in sequence (MCP V1, MCP V2 and penton domain of the spike protein), the triangulation is referred to as ‘pseudo’. Furthermore, as these 21 building blocks are not related by icosahedral symmetry, they all exhibit a slightly different, or ‘quasi-equivalent’, environment, which in theory should be manifested as structural differences^25^. The overall structures of the individual chains were, however, highly similar (Extended Data Figure 5a–c), showing a mean RMSD value of 0.39±0.10 Å (monomers from trimers 1, 2 and 4 relative to a monomer from trimer 3 with three identical chains). Notably, the structural motifs creating intra-trimer contacts (FG1-loop, FG2-loop and FG2-α) showed very little variation (Extended Data Figure 5b). Interestingly, most variation existed in the EF1 and GH1 loops (Extended Data Figure 5c). These loops create inter–trimer contacts, thus acting as molecular switches, and adopt slightly different conformations to accommodate each trimer in their local environment. This is the case especially for loop EF1 which displayed three discrete conformations named ‘outward’, ‘middle’, and ‘inward’ (Extended Data Figure 5d). The outward-facing EF1-loop is used by trimer 1 to reach under the spike penton domain, and analogously by trimer 2 to reach under trimer 1 (Extended Data Figure 5e,f). The inward-facing EF1-loops, in turn, exist solely at the edges of the icosahedral face where the trimers meet the differently tilted trimers of the adjacent face. Finally, the rest of the trimer–trimer interactions are mediated by EF1-loops in the middle position. These observations suggest that a simple structural element alone, such as a loop between two β-strands exhibiting only a few discrete conformational states, can provide an elegant solution for the ‘quasi-equivalence’ problem in the capsid assembly.

**Figure 5.**
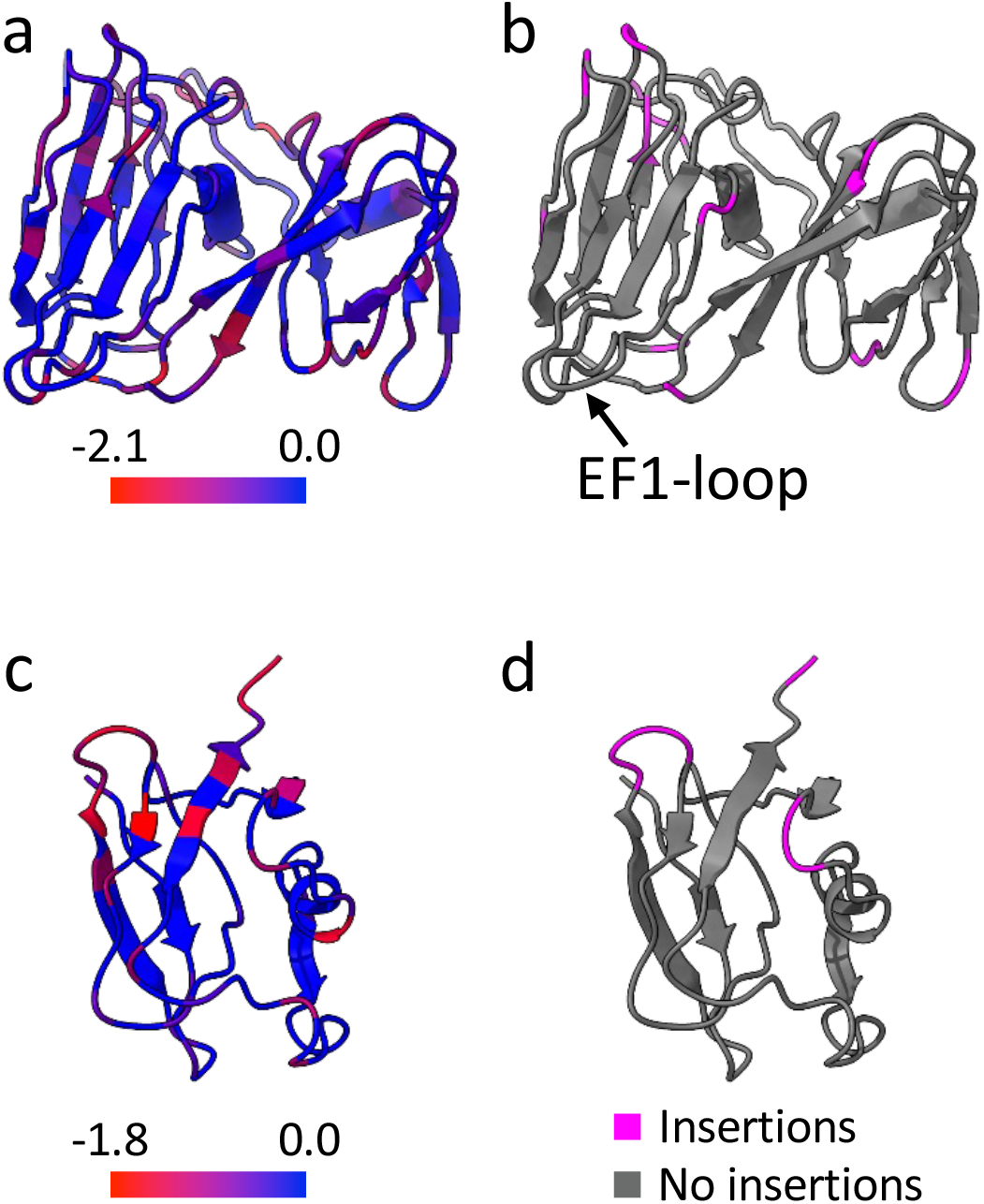
Mapping of ΦCjT23-like prophage sequence variation. **(a)** Sequence conservation^71^ (score –2.12–2.26) determined by multiple sequence alignment of ΦCjT23-like prophage sequences (N=21) and ΦCjT23 major capsid protein (MCP) mapped per residue on the ΦCjT23 MCP structure. **(b)** The same view as in *a* with residues surrounding insertions are highlighted in pink. **(c)** Sequence conservation (score –1.84–2.36) determined by multiple sequence alignment of ΦCjT23-like prophage sequences (N=8) and ΦCjT23 spike protein N-terminal domain. **(d)** The same view as in *c* with residues surrounding insertions highlighted.

### Comparative analysis highlights the minimalistic nature of ΦCjT23 MCP

The MCP (P5) of ΦCjT23 contains only 239 residues, making it the smallest known MCP with a double-jelly-roll fold. The second smallest MCP (269 residues) is that of *Pseudoalteromonas* phage PM2, considered to be the prototypic member of the *Bamfordvirae*^12^. To compare the tertiary structure of the ΦCjT23 MCP to all known double-jelly-roll MCP structures, we constructed a structure-based phylogeny (Fig. 3)^26^. ΦCjT23 MCP, in addition to MCPs of FLiP, PM2, and PRD1, lacks the elaborate decorations created by long loops and other structural elements on top of most other MCPs. The mean of RMSDs between the common core of the ΦCjT23 P5 fold (183 residues) and all analyzed MCPs was 4.4 Å. The closest structural homolog was FLiP MCP with RMSD of 3.0 Å. This phylogeny agrees with the genome type of these viruses as ΦCjT23 and FLiP are the only known ssDNA viruses similar to the members of *Bamfordvirae*, which currently consist exclusively of dsDNA viruses.

In all members of *Bamfordvirae*, the cores of the two β-sandwiches (V1 and V2) in the MCPs are similar, indicative of a gene duplication^4,8,12^. This similarity is especially evident in PM2, ΦCjT23 and FLiP, where the structural alignment of V1 and V2 reports RMSD of 2.0 Å, 2.3 Å and 3.4 Å, over 116, 88 and 105 matching residues, respectively. Further examination of the individual β-sandwich structures and their topologies highlights the minimalistic nature of ΦCjT23 MCP (Extended Data Figure 4). Even the simplest double-jelly-roll MCPs reported earlier show some divergence between V1 and V2 in terms of additional structural elements. For instance, the MCPs of PM2, FLiP, and PRD1 have acquired an additional α-helix, which is membrane-proximal, but can locate either in the N-terminus (PRD1), in a membrane-facing loop (GH-loop of V1 in PM2), or in the C-terminus (FLiP). In PRD1 this helix, in addition to the C-terminus, has been reported to act as a conformational switch. Such an additional helix is absent in ΦCjT23, where the EF1-loop seems to perform the switching function alone. Taken together, these results suggest that V1 and V2 domains of ΦCjT23 have diverged relatively little since the presumed gene duplication^4,8,12^ and ΦCjT23 MCP may thus serve as a model for an ancestral MCP present in the last common ancestor of ΦCjT23 and members of *Bamfordvirae*.

### The ΦCjT23 pentameric viral spike has a unique architecture

The clustering of ΦCjT23 and FLiP MCPs in the structural phylogeny prompted us to investigate whether the structural homology also extends to the vertex complexes of these viruses. First, we analyzed the structure of the ΦCjT23 spike. Localized reconstruction^23^ combined with symmetry relaxation^27^ allowed partial tackling with the observed high degree of spike flexibility by determining the orientations of individual spikes relative to the capsids (Fig. 4a; see Methods). This led to a 5.3-Å resolution structure of the capsid external part of the spike (Fig. 4a,b). Similar to the previously published low-resolution structure of the FLiP spike^13^, the ΦCjT23 spike has a five-fold symmetry (Fig. 4b). At the resolutions reached so far, the protein folds remained elusive, impeding a more detailed comparison. The dimensions and shapes of the two spikes, however, appear different (Fig. 4c)^13^.

In the absence of atomic models for the capsid-external parts of the spikes, we turned our focus to their capsid-internal parts instead. Following the approach applied to the MCPs, we calculated a localized reconstruction of the internal part of ΦCjT23 spike (penton domain), in addition to the FLiP penton base from our previously published data^13^. The resolutions of the localized reconstructions for ΦCjT23 and FLiP were 3.4 Å and 4.0 Å, respectively (Extended Data Figure 6). This was sufficient to build the corresponding atomic models *de novo*. As expected, the structure of the FLiP penton revealed a pentamer formed by five copies of protein P12 (encoded by *orf12*) with the canonical β-sandwich fold, similar to penton base proteins of PM2 and PRD1 (Extended Data Figure 7). This confirmed that the FLiP spike is formed by a separate protein forming a pentamer sitting atop the capsid integral penton base.

**Figure 6.**
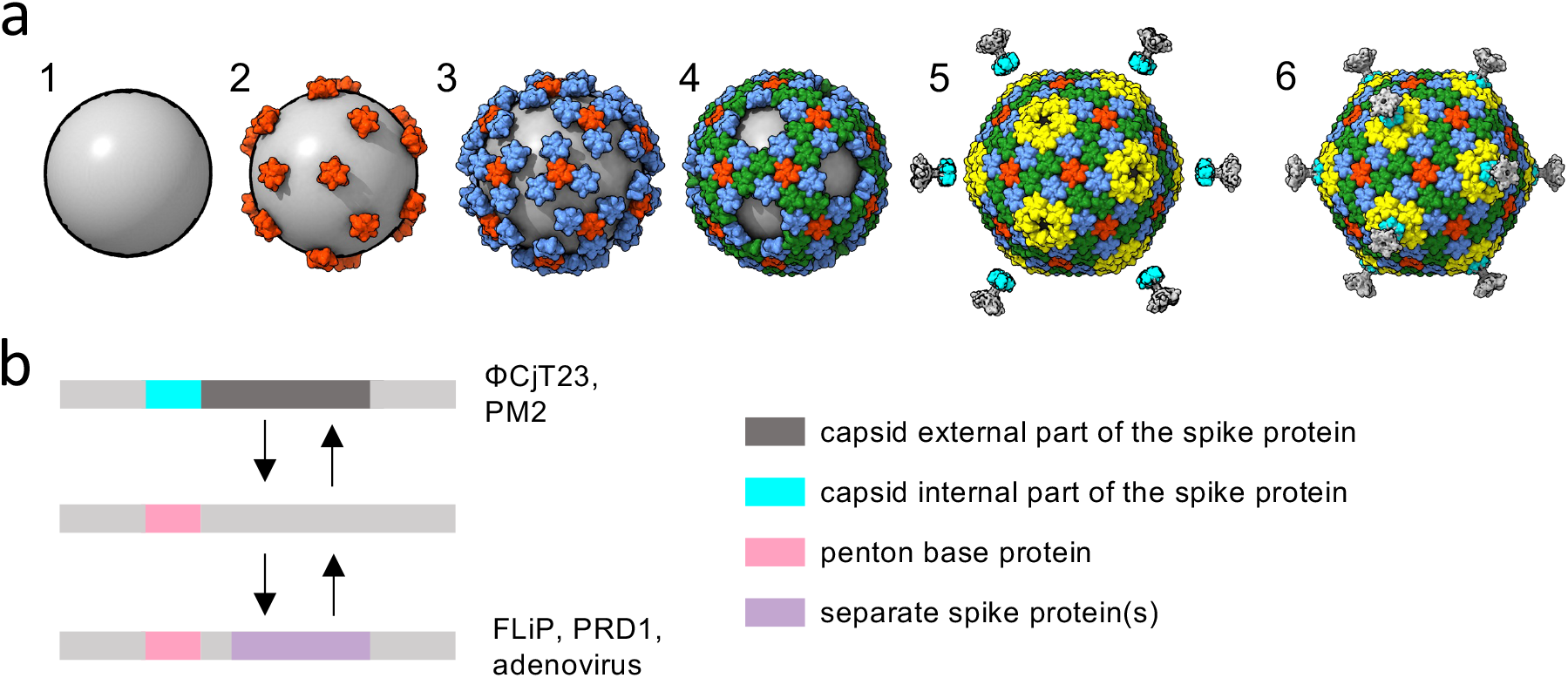
Models for capsid assembly and spike swapping. **(a)** A hypothetical model for the assembly of ΦCjT23 and PM2-like viruses with *T*=21 capsids. First, a membrane vesicle wrapping around the genome assembles to create a lipid core (1). Major capsid protein (MCP) trimers attach to the lipid core via minor membrane proteins (2). More MCPs can be incorporated in a symmetrical fashion via MCP–MCP interactions (3,4) until a nearly closed shell is formed (5). Finally, spikes are incorporated into the capsid (6). **(b)** The genome region encoding for the spikes is depicted in gray. Genetic changes (insertions and deletions) resulting in swapping first the complete spike (such as those observed in ΦCjT23 and PM2) with a penton base and then additional spike proteins being incorporated (such as those observed in FLiP and adenovirus) are depicted.

In contrast to FLiP, the entire spike of ΦCjT23 is made by one protein, P13 (283 residues) (Fig. 1), which forms both the capsid-internal and capsid-external parts, analogously to the spike protein of phage PM2^12^. The capsid-integral part of the spike, corresponding to the P13 N-terminal penton base domain (residues M1–F91), sits in a slightly raised position relative to the type 1 MCP trimers circling it (Fig. 4c). The last residues visible in the cryo-EM density are P90 and F91, residing just prior to the disordered part in the map (Fig. 4d). The sequence immediately following P90 and F91 contains two glycine residues and constitutes a flexible hinge that allows a range of orientations of the external part of the spike relative to the capsid. The penton base domain is a small β-barrel (SBB) with a β-strand topology that differs from the topologies of better-characterized SBBs^28^, and from the canonical β-sandwich fold observed in pentons of other viruses such as FLiP and *Bamfordvirae* members. The ΦCjT23 spike SBB is formed by anti-parallel β-strands β1, β2 and β4, and additional β-strands β3, β5 and β6. The fold encloses a hydrophobic core, characteristic of SBBs^28^. The region connecting strands β5 and β6 includes a short α-helix (P57–I62) and a long loop (N63–E84). The α-helix resides at the monomer– monomer interface. This is analogous to the MCPs where the α-helices reside between V1 and V2 domains of the monomer, in addition to monomer–monomer interfaces. Part of the loop (G66– V72) faces towards the central cavity of the pentamer. Together the five loops close an oculus in the middle of the pentamer (Fig. 4e). Taken together, these results show that the ΦCjT23 spike has a unique architecture with a non-canonical penton domain fold.

What is the significance of the observed differences in the spike architectures of FLiP and ΦCjT23? As the spikes of *Bamfordvirae* members generally mediate host cell recognition^29,30^, we studied this further by testing adsorption and infectivity of ΦCjT23 against three host strains of FLiP (*Flavobacterium sp*. B330, B114, and B167), in addition to the type strain *F. johnsoniae* UW-101. ΦCjT23 infected strains B114 and B167, but not the other two strains tested (B330 and UW-101). Adsorption assays with ΦCjT23 showed similar adsorption patterns in all the strains (including the non-hosts B330 and UW-101) with approximately 35% of phages adsorbed after 5 minutes and the number remaining constant until the end of the experiment (15 minutes, Extended Data Figure 8). However, adsorption to cells of *Flavobacterium* sp. B114 was not observed. It should be noted that because ΦCjT23 can infect this strain, the adsorption step likely requires different conditions than those used in our experiments. In conclusion, ΦCjT23 can adsorb to geographically distant *Flavobacterium* strains, including some of the host strains of its closest relative FLiP, in addition to infecting some of them.

### Mapping of prophage sequence variation on the ΦCjT23 structures

Are the ΦCjT23-like prophages detected in *Flavobacterium* genomes still able to form capsids or have they accumulated mutations that are unlikely to be compatible with capsid assembly? To address this question, we mapped the sequence variation in the homologous prophage ORFs on the structures of the ΦCjT23 MCP and spike N-terminal penton domain (Fig. 5). In both cases, the secondary structure elements are relatively conserved with the greatest sequence variation localizing to loops (Fig. 5a,c). Furthermore, insertions that were present in some of the prophage sequences mapped exclusively to the loop areas (Fig. 5b,d). A notable exception is the EF1-loop that acts as a conformational switch; little sequence variation and no insertions were observed in this loop (Fig. 5a,b).

The presence of both MCP and spike protein sequences in ΦCjT23-like prophages (Extended Data Figure 1) suggests that these can in principle assemble particles with the same mechanism as ΦCjT23 and the sequence variation present in the prophages is unlikely to impede particle assembly. It is possible that the last common ancestor of ΦCjT23 and FLiP utilized the canonical penton fold, which itself has been proposed to be homologous to the MCP domain^9^, and that ΦCjT23 has lost this structural component. Alternatively, ΦCjT23 may be more similar to an ancestral virus that lacked the canonical penton base. This interpretation is supported by the fact that the ΦCjT23 spike sequence is present in several ΦCjT23-like prophages.

## Discussion

The double-jelly-roll-fold is thought to have originated by gene duplication and trimerization presumably led to the basic, hexagon-shaped building block of a primordial capsid^4,8,12^. What is the simplest possible viral capsid built of such pseudo-hexameric MCPs? Classification of different icosahedrally symmetric capsid types based on their *T*-number allows us to systematically consider different structural constraints (Extended Data Figure 8). From a geometric point of view, one constraint is that the MCP is a trimer and thus cannot occupy the two-fold symmetric position. This rules out several capsid types that would otherwise be possible (*T*=4, 12, 16, 28…). Another constraint is the hexamer complexity (*C*^*h*^)^31^. Capsids with hexameric or pseudo-hexameric building blocks with *C*^*h*^ larger than two (*T*=12, 19, 27…) are underrepresented in the observed capsid structures and have been proposed to be disfavored in evolution. A notable exception is the large satellite virus Sputnik, with a *T*=27 capsid and *C*^*h*^=4^16^. Considering these constraints, the smallest allowed – and possibly favoured – *T*-numbers are thus 3, 7, 9, 13, 21 and 25. Of these, *T*=21 is the smallest that has so far been observed in three cases, namely ΦCjT23 in this study, in addition to FLiP^13^ and PM2^15^, whereas *T*=25 has been observed in PRD1^32^, Bam35^33^, and adenovirus^34^. It is conceivable that among capsids built with a double-jelly-roll fold, *T*=13 and smaller capsids would be too small to enclose a lipid bilayer with physically possible membrane curvature and a genome with sufficient length to encode the two required structural proteins and other factors necessary for infection and assembly. Together, these considerations could explain why *T*=21 is the smallest *T* observed for these capsid types. Increased sampling of the virosphere may discover double-jelly-roll-fold-based capsids with smaller *T* than 21 but it remains to be seen if such capsids could encompass a lipid bilayer.

Summarizing the structure of the prototypic ΦCjT23 and analysis of ΦCjT23-like elements in bacterial genomes allow us to propose a model for capsid assembly and evolutionary mechanisms for spike swapping (Fig. 6). First, a p*T*=21 capsid has a trimer at the three-fold axis of symmetry, an arrangement that allows MCP-driven capsid assembly to proceed from this point in a simple, symmetric fashion (Fig. 6a). Such an assembly pathway would not require minor proteins to modulate the conformation of the MCP or tape-measure proteins to define the dimensions of the capsid faces^9^. Instead, the only prerequisites would be a linkage to the membrane (which in turn provides a scaffold for different faces of the capsid to bind to as well as creates the required curvature to form a closed capsid) and a conformational switch in the MCP itself allowing it to lock in place in different positions. Second, the pentameric spikes are required to fill the five-fold positions via interactions with the peripentonal MCPs. These interactions ensure that the spikes, which presumably carry out essential receptor binding functions, are incorporated into the assembly.

The proposed capsid assembly mechanism applies to both ΦCjT23 and PM2. An alternative scenario, where the assembly starts from the vertices, or a special vertex that is structurally distinct from the rest, is also plausible. However, the unique fold of the ΦCjT23 spike penton domain suggests that this structural element is not central to the assembly of these relatively simple *T*=21 capsids. Other viruses utilizing the double-jelly-roll fold have evolved complex capsid architectures requiring minor proteins and more elaborate spike assemblies. For instance, the spikes of adenovirus^35^ and FLiP, are formed by a protein separate from the penton base. Such separate spikes may be more readily swapped to other proteins than those spikes which are formed of a single protein (Fig. 6b). These viruses may have gained entry to new hosts by this proposed mechanism of spike swapping, thereby increasing their fitness. This notion is supported by the plethora of different spike architectures observed in members of *Bamfordvirae*. PRD1 and its close relative PR772 present even more complex vertex architectures. In these viruses, a trimeric spike binds to the capsid by forming a heteropentameric penton base with a penton protein. A monomeric second spike is attached to this penton^36,37^.

In conclusion, this study, together with our earlier findings on FLiP^13^, solidifies a link between dsDNA and ssDNA viruses with capsids utilizing trimeric building blocks with the double-jelly-roll fold, a hallmark of the *Bamfordvirae* viral kingdom. No structures of RNA viruses with similar capsid protein folds have been reported so far. It is thus possible that the last common ancestor of ΦCjT23 and members of the *Bamfordvirae* arose from DNA-based genetic elements, separately from RNA viruses, and co-evolved with cellular life leading to increasingly elaborate variations of the same basic structural principles. Both phages FLiP and ΦCjT23 parasitize *Flavobacterium* species, which are commonly found in both aquatic and terrestrial environments^38-40^ as well as in ancient permafrost^41^ and glacier samples^42^. The presence of these phage types as prophages is in line with their global distribution and common evolutionary origin. Furthermore, our empirical data show that ΦCjT23 adsorbs to several *Flavobacterium* strains, some of which are hosts to FLiP. As ssDNA phages are more prevalent in the environment than often acknowledged^43-45^, further studies are required to establish if similar types of membrane-containing ssDNA phages infect members of other bacterial genera. Understanding of the minimal requirements for capsid assembly from the simplest possible structural elements in this group of viruses may open avenues for engineering of membrane-containing virus-like particles for different applications.

## Methods

### Bacteria and phage

Bacterial strain *Flavobacterium johnsoniae* HER1326 (UW101-36) and phage ΦCjT23 (HER326) were obtained from the Félix d’Hérelle Reference Center for Bacterial Viruses (www.phage.ulaval.ca). *Flavobacterium* sp strain B183 was originally isolated from a freshwater sample in Finland^46^. Bacteria were grown at 25 °C in constant agitation at 150 rpm while high tryptone cytophaga (HTC) medium and 20% Lysogenic Broth (LB) were used as medium. For phage propagation, HTC medium was supplemented with 3 mM MgCl_2_ and 10 mM CaCl_2_. For solid medium, 1 % agar was used while 0.5% Geltrite was used when phage plaque formation was required. The host range of ΦCjT23 was tested against the known hosts of the ssDNA phage FLiP (*Flavobacterium* sp strains B114, B330 and B167) in addition to *F. johnsonia*e UW101.

### Phage adsorption assay

Adsorption of ΦCjT23 to the cells of five *Flavobacterium* strains (the host strain HER1326, *F. johnsoniae* UW101 and *Flavobacterium* sp strains B114, B167 and B330) was tested by adapting an earlier method^47^. In brief, bacteria were grown to mid-log phase and diluted to OD_595_ ∼0.1 with HTC (supplemented with 3 mM MgCl_2_ and 10 mM CaCl_2_). These were divided into triplicates (9 ml in 125 ml Erlenmayer flasks) and 1 ml of ΦCjT23 (∼2.4x10^5^ PFU) was added. Growth medium without bacteria was used as control. Samples of 50 μl were taken into pre-chilled tubes containing 950 μl medium and filtered (0.45 μm). From HER1326 samples were taken at time points 2, 5, 7, 10 and 15 minutes and from others at 5, 10 and 15 minutes and 100 μl of sample was plated with 500 μl of ∼1/5 diluted HER1326 on HTC Gelrite plates.

### Purification of virions

Phages were propagated by transferring the o/n grown bacterial host HER1326 to a fresh HTC medium (supplemented with 3 mM MgCl_2_ and 10 mM CaCl_2_) at a 1:10 ratio and adding a fresh lysate of ΦCjT23 at ratios between 1:100 and 1:200. After six hours of incubation (25 °C, 150 rpm), cells were pelleted by centrifugation (Sorvall RC-6+, F9 6x1000 rotor, 5,400x g, 30 min, 4 °C) and the supernatant was filtered through bottle top filter (0.22 um, Nalgene). Phages were precipitated by dissolving NaCl (0.5 M) and PEG6000 (10 %) to the supernatant and pelleting (Sorvall RC-6+, F9 6x1000 rotor, 5,400× g, 45 min, 4 °C). The resulting pellet was dissolved in 0.02 M Tris-HCl, 3 mM MgCl_2_, and 10 mM CaCl2 pH 7.2. Rate zonal and equilibrium centrifugations were used for subsequent purifications. First, phage suspension was layered on top of a 5–20 % sucrose (w/vol, in Tris-HCl pH 7.2 with 3 mM MgCl_2_ and 10 mM CaCl_2_) and centrifuged (Beckman Optima L-K90, SW-28 rotor, 80,000× g, 45 min, 15 °C). The light-scattering zone was collected and further purified in 20– 70 % sucrose gradient (w/vol, in Tris-HCl pH 7.2 with 3 mM MgCl_2_ and 10 mM CaCl_2_) by centrifugation (Beckman Optima L-K90, SW-41 rotor, 111,000× g for 18 h, 4 °C). The light-scattering zones were collected and pelleted by centrifugation (Beckman Optima L-K90, 70 Ti rotor, 76,000× g for 3 h, 4 °C).

### Analysis of purified virions

Titers (PFU/ml) of the phage suspensions after each purification step were determined. The protein concentrations of the pelleted light-scattering bands from rate zonal and equilibrium centrifugations were determined using the Bradford assay^48^. Structural proteins of the purified virions from the rate zonal centrifugation were resolved by 17 % Tricine SDS-PAGE. The gel was first dyed with Coomassie brilliant blue to detect the proteins after which Sudan Black dye was used for indicating putative lipid structures. Purified virions of phage PRD1 were used for comparison.

### Phage genome isolation, sequencing, and bioinformatics

Phage genomic DNA was extracted from a filtered phage lysate using a ZnCl_2_ precipitation method^49^ with slight modifications. The complementary strand was synthesized using hexamer primers and Klenow fragment. Sequencing was done with Illumina MiSeq sequencer using paired-end (600 bp) kit at the Institute of Biotechnology, University of Helsinki. The assembled genome was further sequenced using Sanger sequencing to confirm uncertain bases. Putative open reading frames were predicted using Glimmer^50^ and GeneMarkS^51^. The NCBI programs BLASTp and DELTA-BLAST against the nonredundant GenBank protein database were used for identifying homologous proteins to the translated ORF sequences^52^. In addition, HHPred search^53^ was employed also for predictions of transmembrane segments.

### Bacterial genome sequencing and analysis

*Flavobacterium* sp B183 Genomic DNA was purified using a QIAGEN Genomic tip 20/G kit, according to manufacturer instructions. Single molecule real-time sequencing was performed on a PacBio RSII sequencer (Génome Québec Innovation Centre) as described previously^54^.

### Prophage identification

BLASTp search for most of the open reading frames predicted in ΦCjT23 genome received several hits to *Flavobacterium* genomes. The resulting hits and their surrounding genome regions were investigated in more detail. In the case of hits to both replication initiation factor containing protein and a putative major capsid protein (hits to ORF that received a hit to FLiP MCP in HHPred search) the regions were subjected to a detailed analysis for identification of integration sequences. WebLogo (available at weblogo.berkeley.edu) was used to create sequence logos for the identified conserved areas at both of the prophage genome ends. Selected 38 prophages sequences filling the above-mentioned criteria were aligned using MAFFT^55^. Some of the prophage sequences were similar (above ∼80% identity) thus one sequence was selected as a representative for the group for further analysis (Supplementary Data Table 2). Easyfig^56^ and BLASTp algorithm were used to create a comparison between the prophage sequences (Extended Data Figure 1).

### Cryogenic electron microscopy of virions

Copper grids coated with holey carbon film (Quantifoil Cu 1.2/1.3, 300 mesh) were glow-discharged for 30 s at 0.45 Torr. A 3-μL aliquot of the purified virions was applied to the grids and after 30 s blotted for 4.5 s and plunge-frozen in liquid ethane with a vitrification apparatus (Vitrobot Mark IV, Thermo Fisher Scientific) equilibrated to 6 °C and 100% relative humidity. The grids were imaged using Talos Arctica transmission electron microscope, operated at 200 kV and equipped with a Falcon III direct electron detector (Thermo Fisher Scientific) in counting mode. A total of 1,023 movies were collected (30 frames per movie) with an electron dose of 0.5 e^−^/Å^2^/frame, resulting in a total dose of ∼15 e^−^/Å^2^. A nominal magnification of 120,000× was used, which resulted in a calibrated pixel size of 1.24 Å. A second data set (852 movies) was collected at a 20-degree stage tilt angle using the same data acquisition parameters as for the first dataset. The two datasets were combined for processing. Data acquisition parameters are summarized in Extended Data Table 1.

### Single-particle data processing and refinement

Data processing was performed using RELION 3.0^57^ as part of Scipion software framework (version 2.0)^58^. Movie frames were aligned with MotionCorr^59^. CTF was estimated with GCTF^60^ (first data set) and CTFFIND4^61^ (second data set). The defocus values ranged from 0.1 μm to 2.2 μm. The particles were picked with ETHAN picker^62^, using a search radius of 240 pixels (first data set) or CrYOLO^63^ using a manually picked set of 301 particles as training data (second data set). Particles (total of 8,862) were extracted in a box of 800×800 pixels. A subset of particles (6,501) was selected for further processing after 2D and 3D classification. The previously determined structure of the FLiP virion (EMD-3771), filtered to 25 Å resolution to reduce template bias, was used as an initial volume. The final map was reconstructed by running RELION’s Bayesian particle polishing, CTF refinement (per-particle defocus and beam tilt) and gold standard 3D auto-refinement applying icosahedral symmetry. A mask defining the capsid shell with a 12-pixel soft edge was created and the resolution was estimated using Fourier shell correlation (FSC) with 0.143 threshold.

### Localized reconstructions of the major capsid protein trimers

To further improve the resolution of the icosahedrally symmetric reconstruction of the capsid, the localized reconstruction method was applied on the four different types of MCP trimers (1–4) to take into consideration the defocus gradient and possible flexibility of the capsid^23,64^. The positions of the trimers were manually defined in UCSF Chimera by placing a marker in the center of each trimer in one asymmetric unit. A total of 390,060 sub-particles (60 for each capsid) were created for trimers 1–3. Trimer 4 position, which is on the icosahedral 3-fold axis of symmetry, yielded 130,020 sub-particles (20 for each capsid). The sub-particles were aligned by placing the local three-fold symmetry axis of each trimer on the Z-axis. Sub-particles were extracted in a box of 200×200 pixels and subjected to one round of 3D classification without alignment, before gold standard, local 3D auto-refinement in RELION. In local refinement, shifts of ±3 pixels were considered with an initial sampling of 1 pixel. Initial angular sampling in the local alignment was 0.9 degrees. Three-fold symmetry was applied on hexamer 4. No symmetry was applied on trimers 1–3. A mask with a 3-pixel soft edge was created for each trimer localized reconstruction and the resolution was estimated as above.

### Localized reconstruction of the penton

Similar to the trimer localized reconstructions, we separately reconstructed the penton base. A total of 78,012 penton sub-particles (12 for each capsid) were defined, aligned in C5 symmetry convention, extracted, and refined as above. Five-fold symmetry was applied. The map was masked, and its resolution was estimated as above.

### Localized reconstruction of the spike

To facilitate localized reconstruction of the spikes, the contribution of the icosahedral capsid was subtracted from the particle images. A mask defining the capsid and its interior was created by a thresholding operation, applied on a lowpass filtered (15 Å) version of the capsid reconstruction, followed by dilation and erode operations to fill holes in the mask. Partial signal subtraction of the capsid was performed in RELION^65^ considering the CTF of each particle. Sub-particles (78,012) were defined at a radius of 315 Å as above and extracted from the capsid-subtracted images. To find the tilt direction of each spike, particles were aligned using “*relax_sym C5*”^27^ option in RELION 3D classification. Only the five directions related by C5 symmetry were considered. Other angular and translational parameters were fixed. No symmetry was applied, and two classes were used. Spikes in one class were severely tilted and excluded from further processing whereas spikes in the other class were only moderately tilted and were included. To reconstruct an initial map of the moderately tilted spike, RELION 3D auto-refinement was used with 3.7 degrees initial angular sampling and default search range. No symmetry was applied. To re-align the spike C5 symmetry axis, another local alignment was carried out using 7.5 degrees initial angular sampling and custom angular search range sigma of 10 degrees (option *sigma_ang*). To compensate for possibly large rotations of the spike, shifts of ±10 pixels were allowed. To calculate the final reconstruction of the spike, the sub-particle alignment parameters were adjusted by applying the required rotation to align their C5 symmetry axis on the Z-axis. This was performed using protocol ‘*xmipp – align volume and particles*’ and volume alignment of the initial asymmetric spike reconstruction on the initial C5 spike reconstruction. The adjusted spike orientations were considered as priors in the final round of RELION local auto-refinement using 3.7 degrees initial angular sampling and default search range. This workflow improved the resolution of the spike from 7.2 Å (tilted spike, no symmetry) to 5.3 Å (aligned spike, C5 symmetry) and B-factor from 420 Å^2^ to 219 Å^2^, respectively.

### Composite model of the capsid

Protocol ‘*localrec – stitch subvolumes*’ in Scipion plugin LocalRec 3.0.2 was used to combine the localized reconstructions of trimers 1–4, penton base and the spike into a composite map of the icosahedral capsid^64^. A spherical mask with a diameter of 80 pixels was used for the trimers and the penton. To define the boundary between the high-resolution penton map and the low-resolution spike map, a spherical mask defining the spike (diameter 100 pixels) was first subtracted from the spherical mask of the penton after an appropriate shift reflecting the different centering of the two localized reconstructions relative to the capsid. This ensured that signals for the shaft and linker regions of the spike were maximally retained in the composite map. Prior to running the protocol, the density values in the low-resolution spike map were scaled by a factor of two which was determined by trial and error to allow their visualization as a single isosurface.

### Localized reconstruction of the FLiP penton

Localized reconstruction of the FLiP penton was calculated similar to the ΦCjT23 penton (see above). Particles (2,351) were extracted from a previously published data set of 235 micrographs. A total of 28,212 penton sub-particles were defined at a radius of 300 Å and extracted in a box of 200×200 pixels. One round of local RELION 3D auto-refinement with an initial angular sampling of 1.8 degrees and a custom search range of 2 degrees (option *sigma_ang*) resulted in a FLiP penton map with a resolution of 4.0 Å. Data acquisition parameters are summarized in Extended Data Table 2.

### Atomic model building and refinement

To identify ORFs encoding ΦCjT23 MCP, penton, and FliP penton, polyalanine models of each target protein were built into their respective cryo-EM reconstructions in COOT^66^. Secondary structure elements found in the polyalanine models were compared to the secondary structure elements predicted for all ΦCjT23 and FLiP ORFs using JPred^67^, and sequences from ORFs with similar profiles were built into the cryo-EM reconstructions in COOT in order to validate matches between sequence and density. Following ORF identification, model building, and structure refinement were performed using COOT and PHENIX^68^ in consecutive cycles with secondary structure and geometry restrained. Finally, the four ΦCjT23 MCP trimer atomic models and the ΦCjT23 penton atomic model were rigid-body fitted back in their original positions of the capsid to create an atomic model of the asymmetric unit of the entire capsid. Refinement statistics of all atomic models are presented in Extended Data Table 1. All figures were rendered in UCSF ChimeraX^69^ and topology diagrams were generated with PDBsum^70^.

### Structural phylogeny

Structural alignments of ΦCjT23 MCP and other viral MCPs with a double β--sandwich fold and a phylogenetic tree were calculated using Homologous Structure Finder software^26^.

## Author contributions

Conceptualization, E.L., S.M., L-R.S., J.T.H.; Formal Analysis, N.K.; Investigation, N.K., E.L., I.R., M.S., J.R., J.T.H.; Writing – Original Draft, J.T.H.; Writing – Review & Editing, all authors; Visualization, E.L., I.R., J.R., L-R.S., J.T.H; Resources, S.M., L-R.S., J.T.H.; Software, V.A.; Supervision, L-R.S., J.T.H.

## Acknowledgements

We thank Mark McBride for bacterial strains and comments on the manuscript as well as Denise Tremblay, Benita Löflund, Kati Mäkelä, Ville Hoikkala and Pasi Laurinmäki for technical assistance. This study was supported by the Academy of Finland (342988 to I.R., 321985 to E.L. and 314939 to L-R.S.) and Emil Aaltonen Foundation (to L-R.S.). The facilities and expertise of the HiLIFE cryo-EM unit at the University of Helsinki, a member of Instruct-ERIC Centre Finland, FINStruct, and Biocenter Finland are gratefully acknowledged. The authors wish to acknowledge CSC – IT Center for Science, Finland, for computational resources. S.M. holds the Canada Research Chair in Bacteriophages.

## Data availability

Data that support the findings of this study have been deposited in the GenBank with the accession code ON067806 (ΦCjT23 genomic sequence) and CP097434 (*Flavobacterium* sp B183 whole genome sequence), in the Electron Microscopy Data Bank (EMDB) with the accession codes EMD-15042, EMD-15044, EMD-15045, EMD-15046, EMD-15047, EMD-15048, EMD-15049 EMD-15050, EMD-15051 (cryo-EM density maps) and in the Protein Data Bank (PDB) with the accession codes 7ZZZ, 8A01, 8A02, 8A03, 8A04, 8A05, 8A06 (atomic models). All other data generated or analysed during this study are included in this published article (and its supplementary information files).

