## Extended Data File for "New Vista into Origins of Viruses from a Prototypic ssDNA Phage"

[*] corresponding authors
[#] contributed equally

### Extended Data Tables

**Extended Data Table 1 | ΦCjT23 cryo-EM data collection and processing details.**

| | **Virion** (EMD-15042) (PDB 7ZZZ) | **Trimer 1** EMD-15044) (PDB 8A01) | **Trimer 2** EMD-15045) (PDB 8A02) | **Trimer 3** (EMD-15046) (PDB 8A03) | **Trimer 4** EMD-15047) (PDB 8A04) | **Spike, penton domain** (EMD-15048) (PDB 8A05) | **Spike, external part** (EMD-15049) |
|---|---|---|---|---|---|---|---|
| **Data collection and processing** | | | | | | | |
| Magnification | 120,000× | 120,000× | 120,000× | 120,000× | 120,000× | 120,000× | 120,000× |
| Voltage (kV) | 200 | 200 | 200 | 200 | 200 | 200 | 200 |
| Electron exposure ($e^{-}$/Å$^2$) | 15 | 15 | 15 | 15 | 15 | 15 | 15 |
| Defocus range (μm) | 0.3–2.2 | 0.3–2.2 | 0.3–2.2 | 0.3–2.2 | 0.3–2.2 | 0.3–2.2 | 0.3–2.2 |
| Pixel size (Å) | 1.24 | 1.24 | 1.24 | 1.24 | 1.24 | 1.24 | 1.24 |
| Symmetry imposed | I1 | C1 | C1 | C3 | C1 | C5 | C5 |
| Initial particle images (no.) | 8,862 | 390,060 | 390,060 | 130,020 | 390,060 | 78,012 | 74,668 |
| Final particle images (no.) | 6,501 | 265,790 | 257,197 | 98,641 | 265,085 | 18,122 | 40,397 |
| Map resolution (Å) | 4.1 | 3.2 | 3.2 | 3.2 | 3.2 | 3.4 | 5.3 |
| FSC threshold | 0.143 | 0.143 | 0.143 | 0.143 | 0.413 | 0.143 | 0.143 |
| Map sharpening *B* factor (Å$^2$) | –151 | –141 | –156 | –139 | –150 | –151 | –219 |
| **Refinement & validation** | | | | | | | |
| Model-to-map resolution (Å) | 4.3 | 3.3 | 3.4 | 3.4 | 3.4 | 3.8 | N/A |
| FSC threshold | 0.5 | 0.5 | 0.5 | 0.5 | 0.5 | 0.5 | N/A |
| Model-to-map CC | | | | | | | |
| Main chain | 0.82 | 0.86 | 0.84 | 0.84 | 0.85 | 0.83 | N/A |
| Side chain | 0.81 | 0.83 | 0.82 | 0.82 | 0.83 | 0.81 | N/A |
| Model composition | | | | | | | |
| Non-hydrogen atoms | 19134 | 5511 | 5511 | 5511 | 5511 | 764 | N/A |
| Protein | 19134 | 5511 | 5511 | 5511 | 5511 | 764 | N/A |
| Ligand | 0 | 0 | 0 | 0 | 0 | 0 | N/A |
| Model resolution range (Å) | N/A | 3.2 | 3.2 | 3.2 | 3.2 | 3.4 | N/A |
| *B* factors (Å$^2$) | | | | | | | |
| Protein | N/A | 33.5 | 31.9 | 45.6 | 34.3 | 45.1 | N/A |
| R.m.s. deviations | | | | | | | |
| Bond lengths (Å) | N/A | 0.002 | 0.003 | 0.002 | 0.004 | 0.002 | N/A |
| Bond angles (°) | N/A | 0.439 | 0.469 | 0.437 | 0.599 | 0.424 | N/A |
| Validation | | | | | | | |
| MolProbity score | N/A | 1.25 | 1.21 | 1.21 | 1.21 | 1.01 | N/A |
| Clash score | N/A | 4.75 | 4.31 | 3.41 | 4.31 | 1.99 | N/A |
| Rotamer outliers (%) | N/A | 0 | 0 | 0 | 0 | 0 | N/A |
| Ramachandran plot | | | | | | | |
| Favored (%) | N/A | 98.45 | 98.03 | 97.61 | 98.45 | 97.83 | N/A |
| Allowed (%) | N/A | 1.55 | 1.97 | 2.39 | 1.55 | 2.17 | N/A |
| Outliers (%) | N/A | 0 | 0 | 0 | 0 | 0 | N/A |

**Extended Data Table 2 | FLiP cryo-EM data processing details.**

| | **Penton** (EMD-15051) (PDB 8A06) |
|---|---|
| **Data processing** | |
| Pixel size (Å) | 1.35 |
| Symmetry imposed | C5 |
| Initial particle images (no.) | 141,060 |
| Final particle images (no.) | 28,212 |
| Map resolution (Å) | 4.0 |
| FSC threshold | 0.143 |
| Map sharpening *B* factor (Å$^2$) | –164 |
| **Refinement** | |
| Model-to-map resolution (Å) | 4.3 |
| FSC threshold | 0.5 |
| Model-to-map CC | |
| Main chain | 0.75 |
| Side chain | 0.70 |
| Model composition | |
| Non-hydrogen atoms | 1185 |
| Protein | 1185 |
| Ligands | 0 |
| Model resolution range (Å) | 4.0 |
| *B* factors (Å$^2$) | |
| Protein | 36.2 |
| R.m.s. deviations | |
| Bond lengths (Å) | 0.003 |
| Bond angles (°) | 0.556 |
| Validation | |
| MolProbity score | 1.47 |
| Clash score | 5.94 |
| Rotamer outliers (%) | 0.73 |
| Ramachandran plot | |
| Favored (%) | 97.24 |
| Allowed (%) | 2.76 |
| Outliers (%) | 0 |

### Extended Data Figure Legends

**Extended Data Figure 1 | Comparison of detected phiCjT23-like prophages.** An Easyfig alignment of the representative prophage regions in *Flavobacterium* and *Lacinutrix* sp (*Flavobacteriaceae)* genomes. Genes with putative functions are marked with colors with indicating functions shown in the bottom. Some of the prophage genes received hits in a BlastP search against ssDNA phages FLiP (all hits to FLiP gp16) and Cellulophaga phages phi48:2, phi12:a:1, phi12:2 and phi18:4, and these hits are also indicated with colors in the bottom. In addition, several of the replication proteins showed similarities with the small Cellulophaga sDNA phages. Also, the identified major capsid protein sequence showed similarities with the structural proteins of these phages.

**Extended Data Figure 2 | Cryo-EM and three-dimensional reconstruction of ΦCjT23 bacteriophage and its capsid components. (a)** A representative cryo-EM micrograph of purified ΦCjT23 particles. One smaller particle, possibly representing the lipid core of ΦCjT23, is marked with an asterisk. **(b)** Representative class averages of particles. One class average, representing the putative lipid core, is marked with an asterisk. **(c)** A 5-Å thick central cross-section of the conventional icosahedral reconstruction. DNA and membrane (M) densities are labelled. The inset shows a close-up (3×) of one spike (arrowhead). **(d)** Areas for different localized reconstructions are illustrated with colored transparent spheres on the icosahedral reconstruction (gray). Note that for clarity, the spheres are rendered smaller than the localized reconstructions. The insets show resulting localized reconstructions of the spike (purple) and vertex (cyan) are shown from side. **(e)** The localized reconstructions of the trimers 1–4 are shown from the top. **(f)** A 5-Å thick central section of a composite volume of the particle (EMD-15050), calculated by combining the localized reconstructions. The inset shows a close-up (3×) of one spike (arrowhead) with improved density over the conventional map (*c*, inset). Scale bars, 50 nm.

**Extended Data Figure 3 | Resolution and map quality of the major capsid protein. (a)** Fourier shell correlation plots (FSC), calculated between two half-maps as a function of spatial frequency, are shown for each trimer (1–4) localized reconstruction. In each case, FSC is plotted for the original unmasked half-maps (gray), masked half-maps (blue), and phase-randomized half-maps (red) in which phases were randomized at frequencies higher than 1/7 Å. The phase-randomized FSC drops sharply at the cutoff frequency below the noise threshold (0.143), as expected. The phase-randomization test was used to take the effect of masking on the half-maps into account before calculating the final, corrected FSC curve (black). Good agreement between the masked and corrected curves indicated that masking did not cause overestimation of resolution. In all cases, the corrected curve drops below the noise threshold (FSC=0.143) at 1/3.2 Å indicating a resolution of 3.2 Å in the reconstructions. **(b–d)** Selected parts of the trimer 3 map V1 domain are shown, namely the FG1 α-helix (*b*), β-sheet C-H-E-F (*c*) and β-sheet B-I-D-G (*d*).

**Extended Data Figure 4 | Major capsid protein antiparallel β-sandwich structures.** Different structural elements are shown as topology diagrams for the N-terminal (V1) and C-terminal (V2) β-sandwich with arrows (β-strands), boxes (α-helices) and arcs (linker elements between β-strands and termini). β-strands are labeled B1–I1 in V1 and B2–I2 in V2. Linkers consist of β-turns or longer loops. The approximate length of each linker element is given. Longer linkers contain in some cases additional secondary structure elements. Short secondary structure elements (four residues or shorter) are not shown for clarity. α-helices residing on the membrane proximal side of the β-sandwich in FLiP, PM2 and PRD1 are colored in red. The insets show the corresponding ribbon representation for each domain. The same elements are highlighted in red as in the topological diagrams.

**Extended Data Figure 5 | Comparison of MCPs chains. (a)** All ten major capsid protein chains from the asymmetric unit (three for trimers 1,2 and 4; one for trimer 3) are shown after aligning them. The two β-sandwiches V1 and V2 are labeled. (**b**) The view in *a* is shown after rotating 90 degrees around the horizontal axis as indicated to view the monomer from outside of the capsid. Note that little structural variation exists in the FG1 and FG2 loops (labeled). (**c**) The view in *a* is shown after rotating 90 degrees around the horizontal axis as indicated to view the monomer from inside of the capsid. The two loops showing the greatest degree of structural variation, EF1 and GH1 loop, are labeled. **(d)** A close-up of the EF1 loops is shown for the area indicated in (*c*). The three discrete conformations are labeled (outward, middle, inward). The arc depicts the range of motions this loop can undergo. **(e)** All chains in our asymmetric unit (colored as in Figure 1) are shown, together with their neighbors (gray). The type of each EF1-loop is indicated (outward, O; middle, M; inward I). The demarcation lines between the icosahedral faces are shown with dashed lines.

**(f)** Examples for each EF1-loop type are shown from the inside of the capsid. The trimer the EF1-loop interacts with is shown as a surface.

**Extended Data Figure 6 | Resolution and map quality of the ΦCjT23 penton domain and FLiP penton protein.** **(a)** Fourier shell correlation (FSC) is shown for the ΦCjT23 penton domain localized reconstruction. Different curves are as in Supplementary Figure 3a. The corrected curve drops below the noise threshold (FSC=0.143) at 1/3.4 Å indicating a resolution of 3.4 Å in the reconstruction. **(b–c)** Examples of the ΦCjT23 penton domain cryo-EM density are shown for an α-helix (*b*) and a β-sheet (*c*). **(d)** FSC is shown for the FLiP penton protein localized reconstruction. The corrected curve drops below the noise threshold at 1/4.0 Å indicating a resolution of 4.0 Å in the reconstruction. **(e–f)** Examples of the FLiP penton protein cryo-EM density are shown for an α-helix (*e*) and a β-sheet (*f*).

**Extended Data Figure 7 | The ΦCjT23 penton domain has a distinct fold**. **(a)** The common double β-barrel penton structure, presented by PRD1 penton protein (PDB:1W8X), are shown. **(b)** The same topology is observed in the FLiP penton. **(c)** ΦCjT23 capsid internal part of the spike (penton domain) shows a distinct, unrelated fold. For each penton structure, a ribbon representation, colored rainbow ramped from blue (N-terminus) to red (C-terminus), is shown (left) alongside a topology diagram (right). Secondary structure elements in the topology diagram are presented with arrows (β-strands) and cylinders (α-helices) and are colored as in the respective ribbon representations.

**Extended Data Figure 8 | Adsorption of ΦCjT23 to the cells of *Flavobacterium*.** Dots represent individual values of three replicates, horizontal line their mean, and vertical line standard error.

**Extended Data Figure 9 | Icosahedral virus capsid architectures.** A schematic diagram is shown for different types of icosahedrally symmetric capsids built of 12 pentagons (gray) and $N = 60 \times (T–1) / 6$ hexagons (white). The triangulation (*T*) number of each capsid is defined by two lattice indexes h and k as $T = h^2 + h \times k + k^2$. In the members of the PRD1–adenovirus lineage (*Bamfordvirae*), the allowed hexagon positions are occupied by pseudohexameric trimers, hence the *T*-number is indicated being pseudo (*p*). Different capsid types are classified in class 1, 2 and 3. Only the first members of the series (…) are shown in each class. The lattice axes are illustrated for one capsid in each class with arrows. Hexagons that are located on the icosahedral two-fold axes of symmetry, and that thus cannot be occupied by a pseudohexameric, trimeric major capsid protein (MCP), are colored in red (corresponding capsids are labelled as “not allowed”). For each capsid, the value of hexamer complexity ($C^h$) is given. Those capsids that are geometrically allowed capsids but where $pT$<21 may be too small to encompass the lipid bilayer and genome. Names of the characterized viruses with a certain capsid type ($pT$ = 21, 25, 27) are given. Capsid geometries with no characterized representatives are labelled as unobserved. Note that the lattice in class 2 capsids is handed and only the right-handed (dextro) organization, which is observed in in PM2, FLiP and ΦCjT23 is shown. Allowed capsid geometries ($pT$=3, $p$T=9, $pT$=21 and $pT$=27) where a hexagon occupies the three-fold axes of symmetry are indicated by coloring one such hexagon blue.

### Extended Data Figure 1

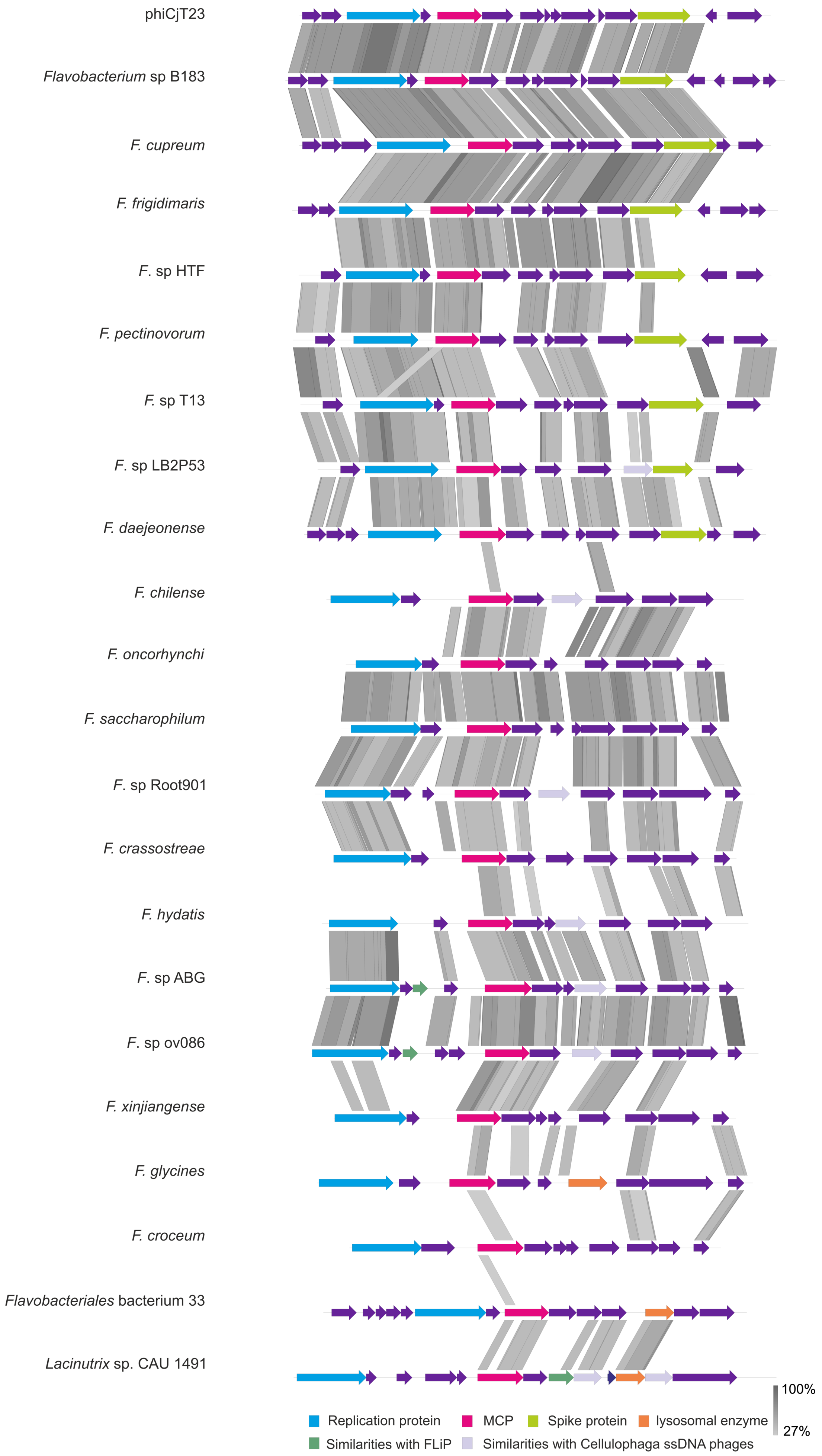

Extended Data Figure 2

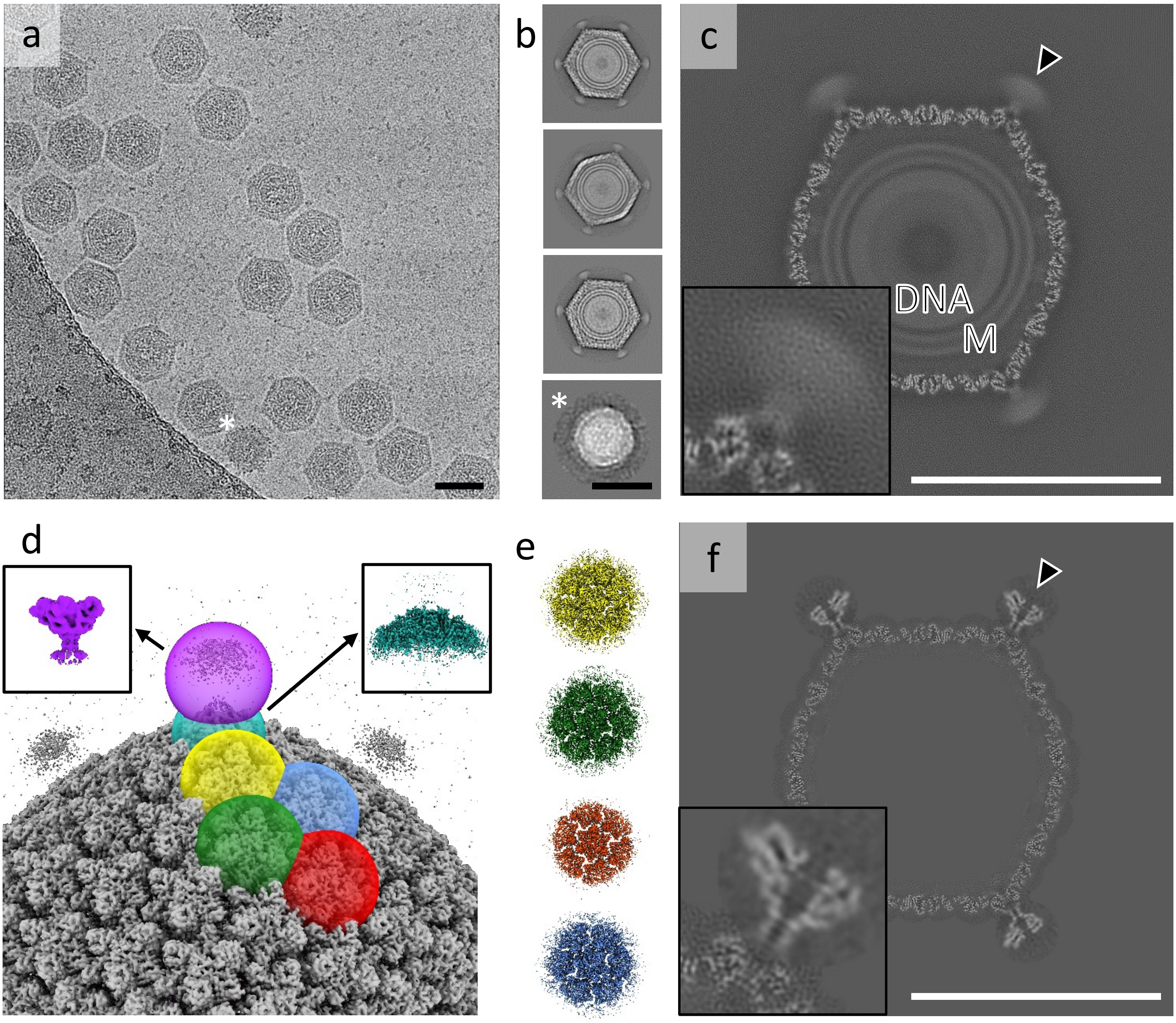

Extended Data Figure 3

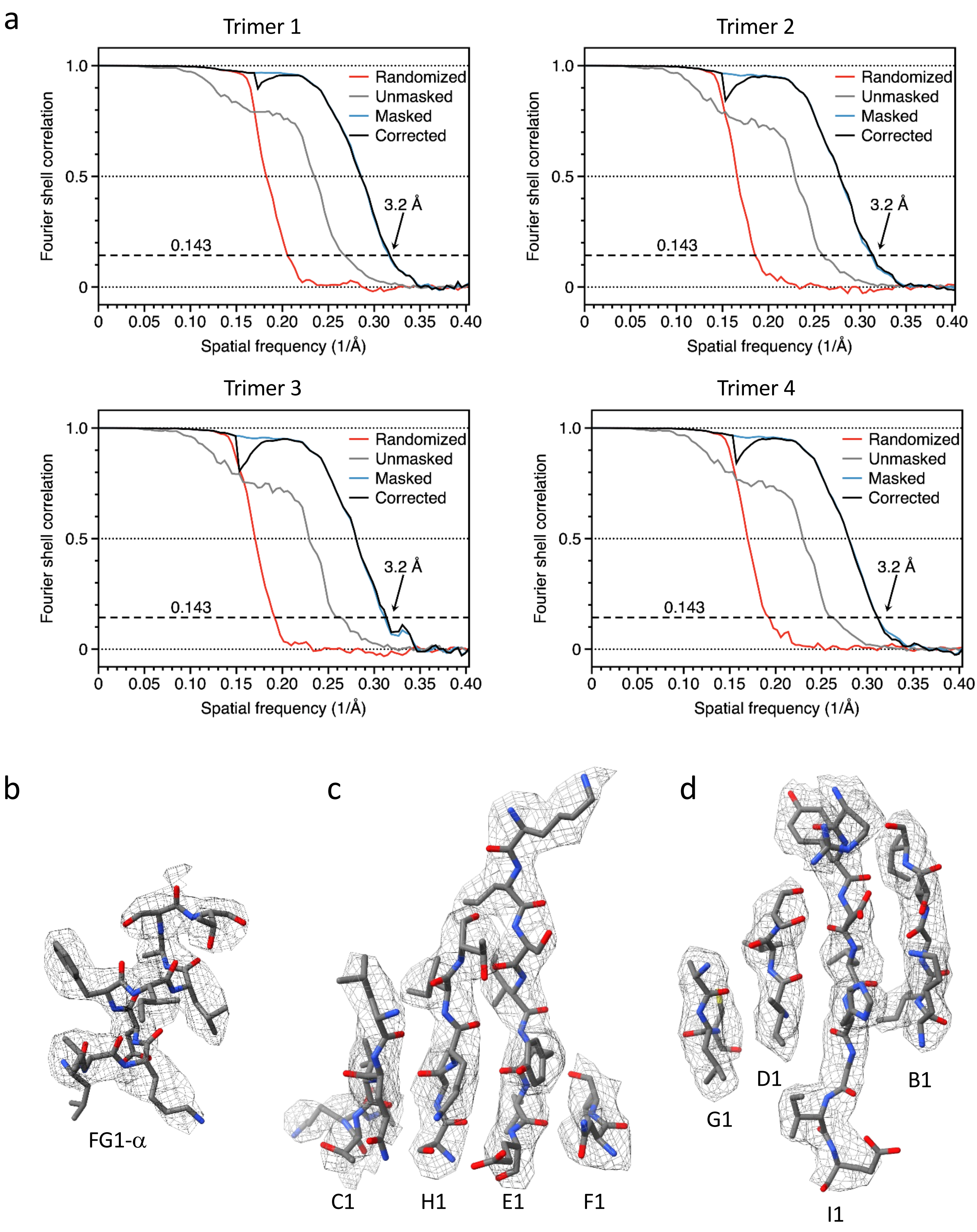

### Extended Data Figure 4

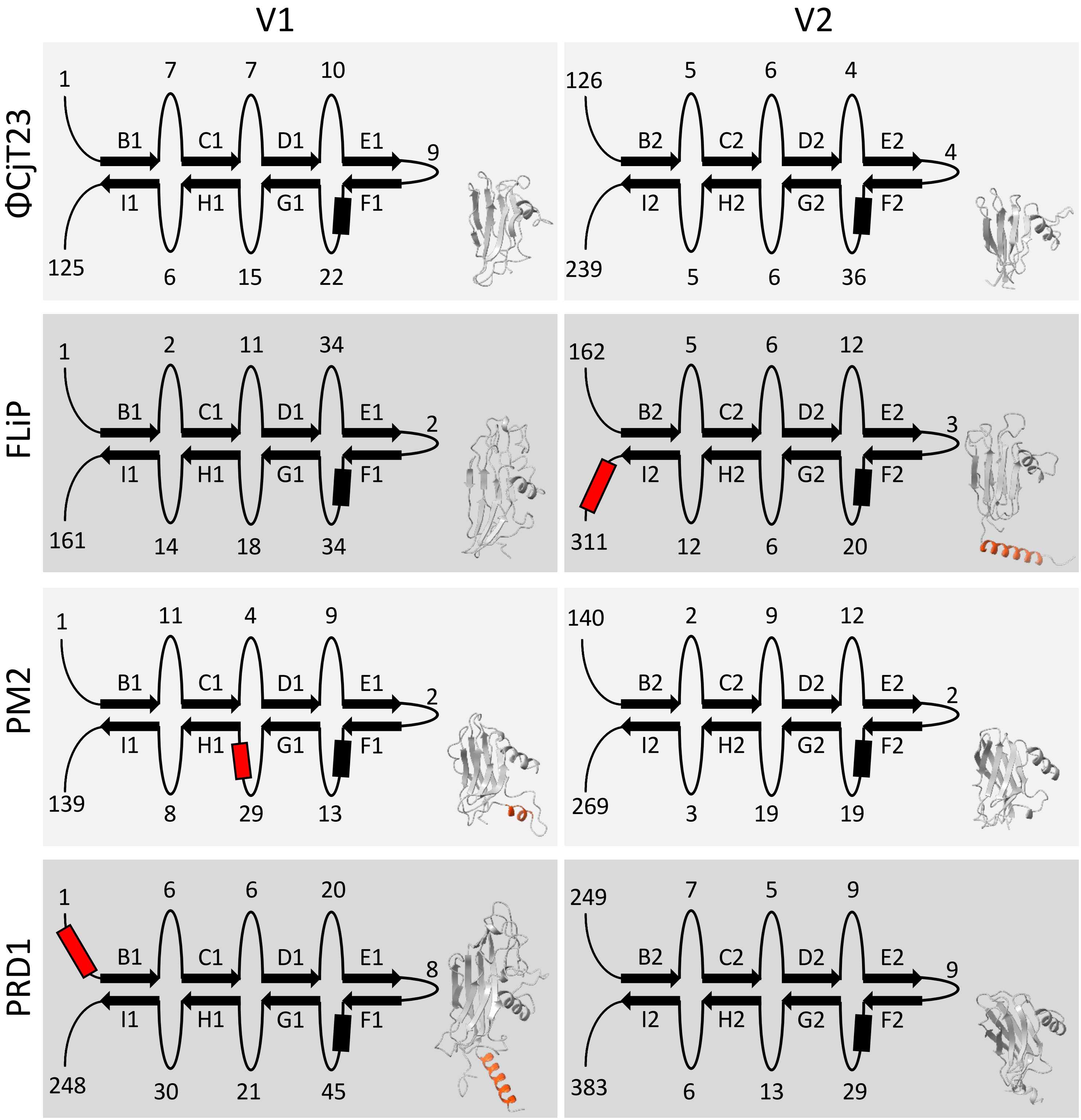

Extended Data Figure 5

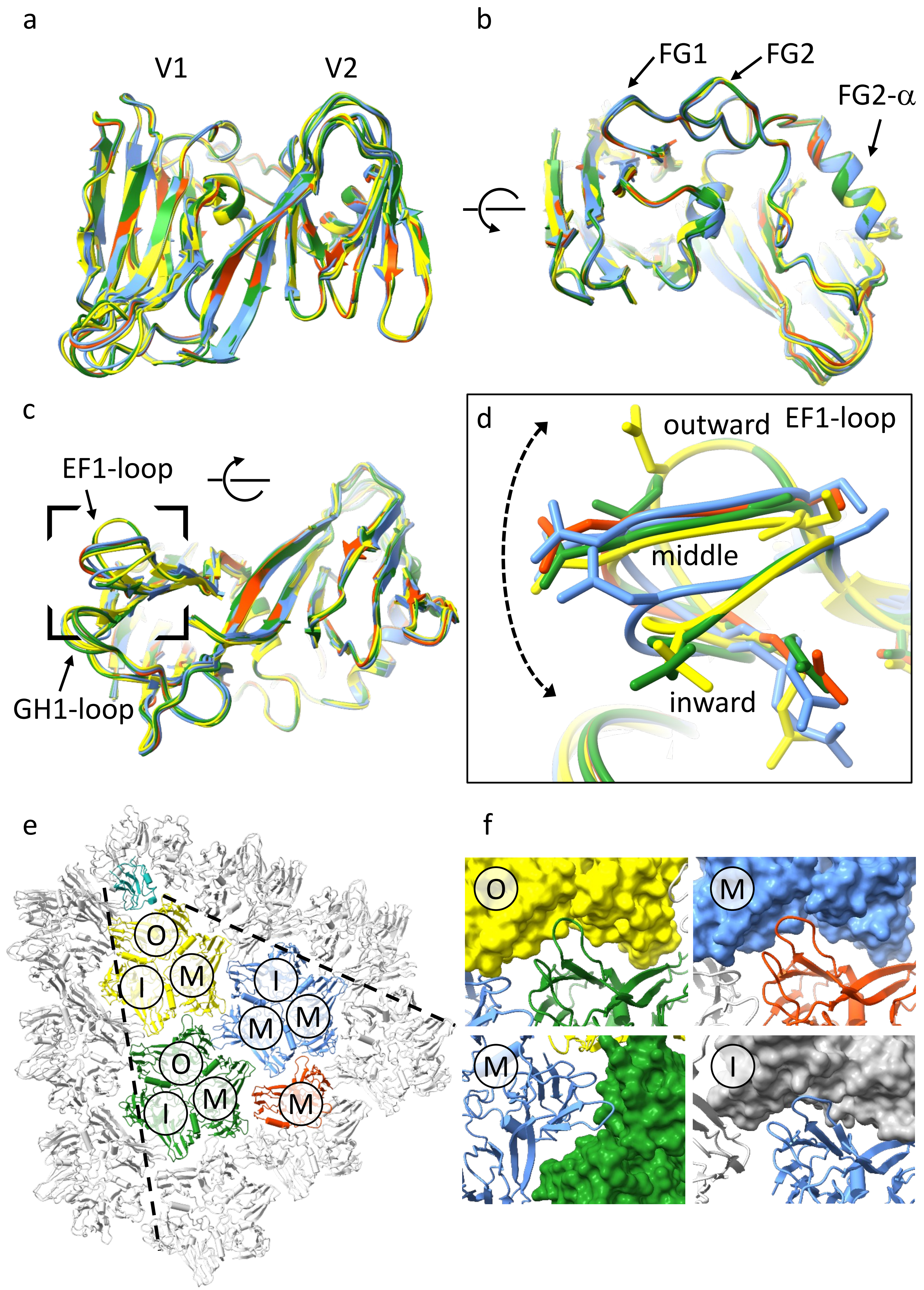

### Extended Data Figure 6

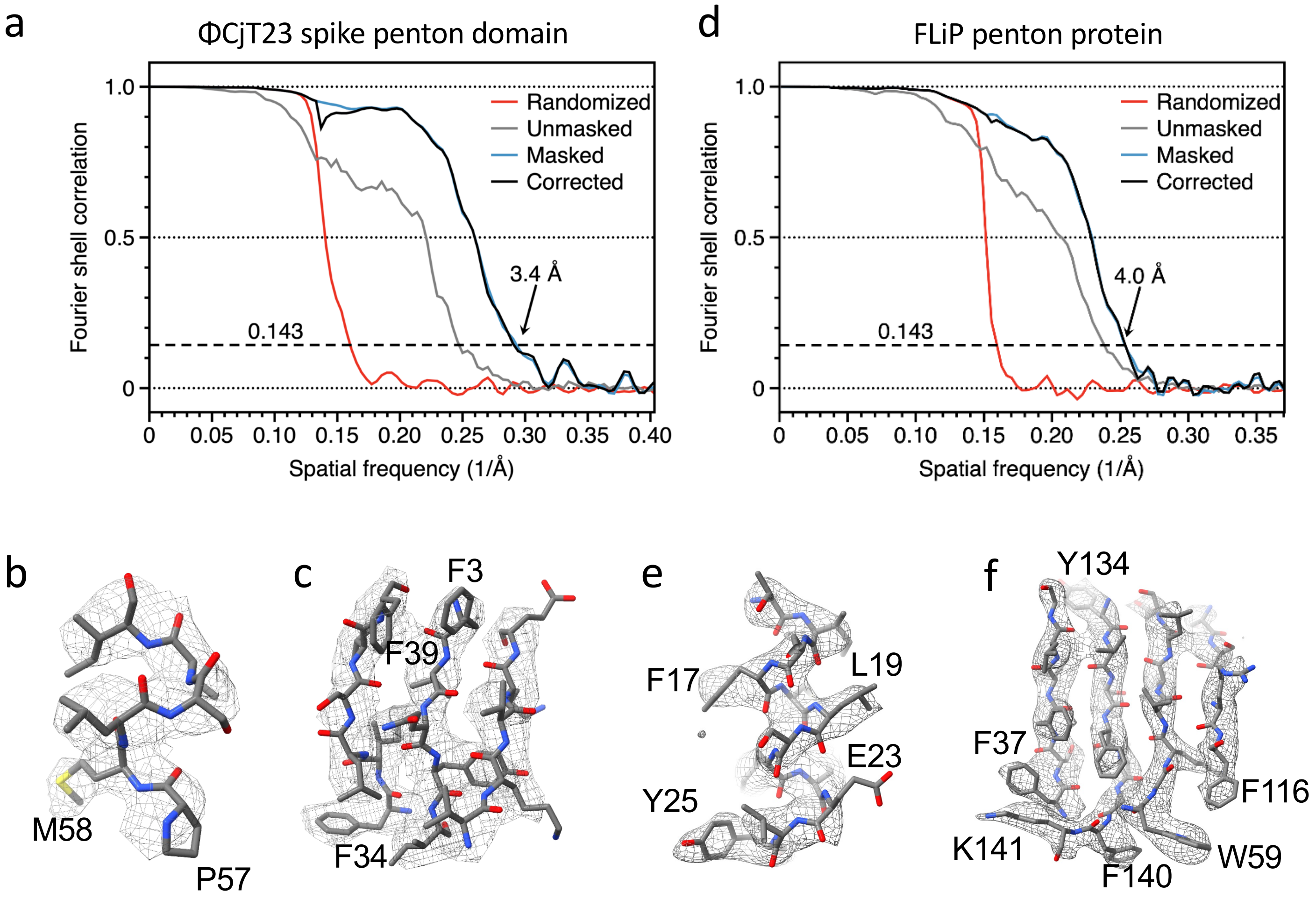

### Extended Data Figure 7

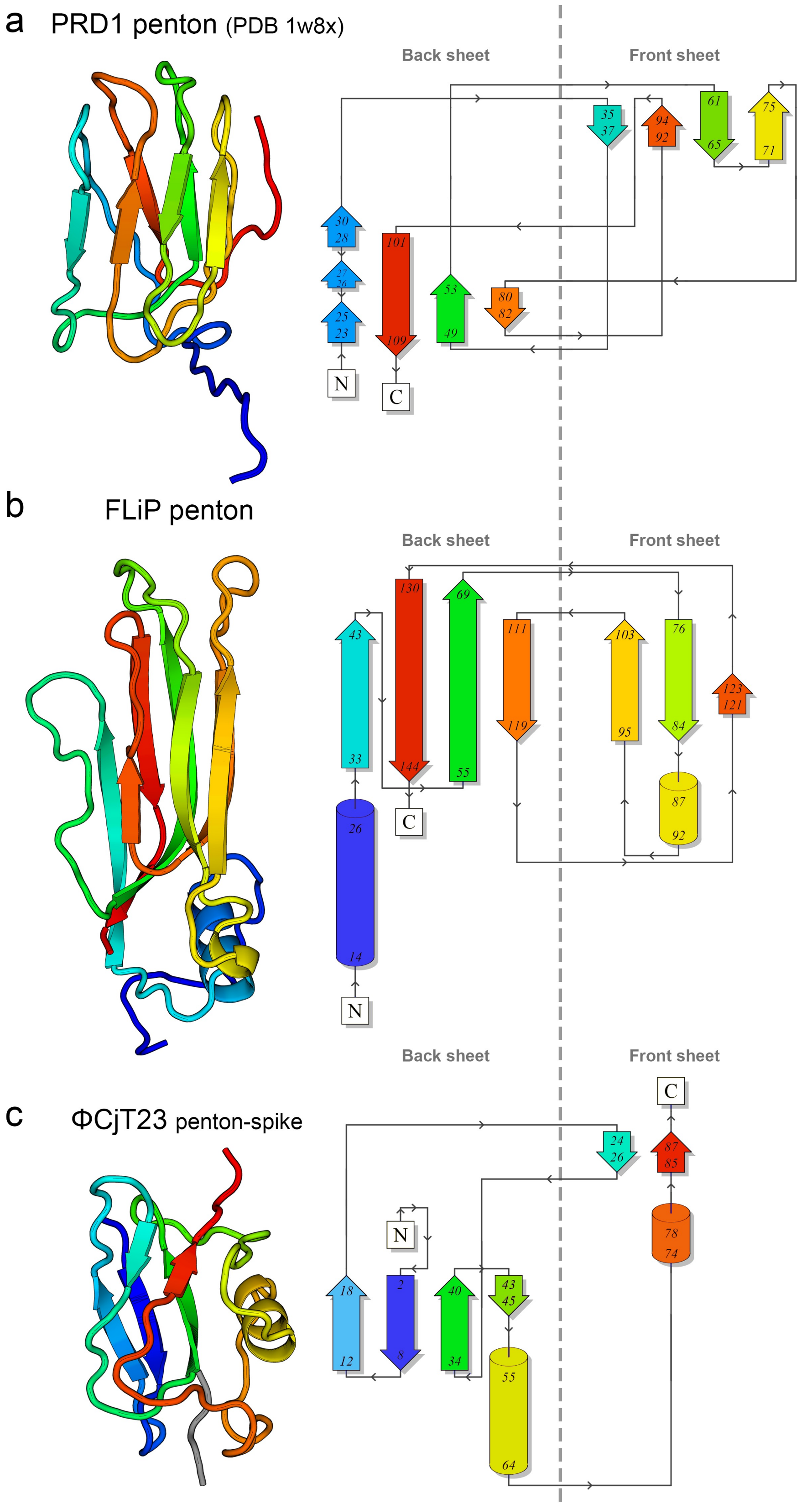

### Extended Data Figure 8

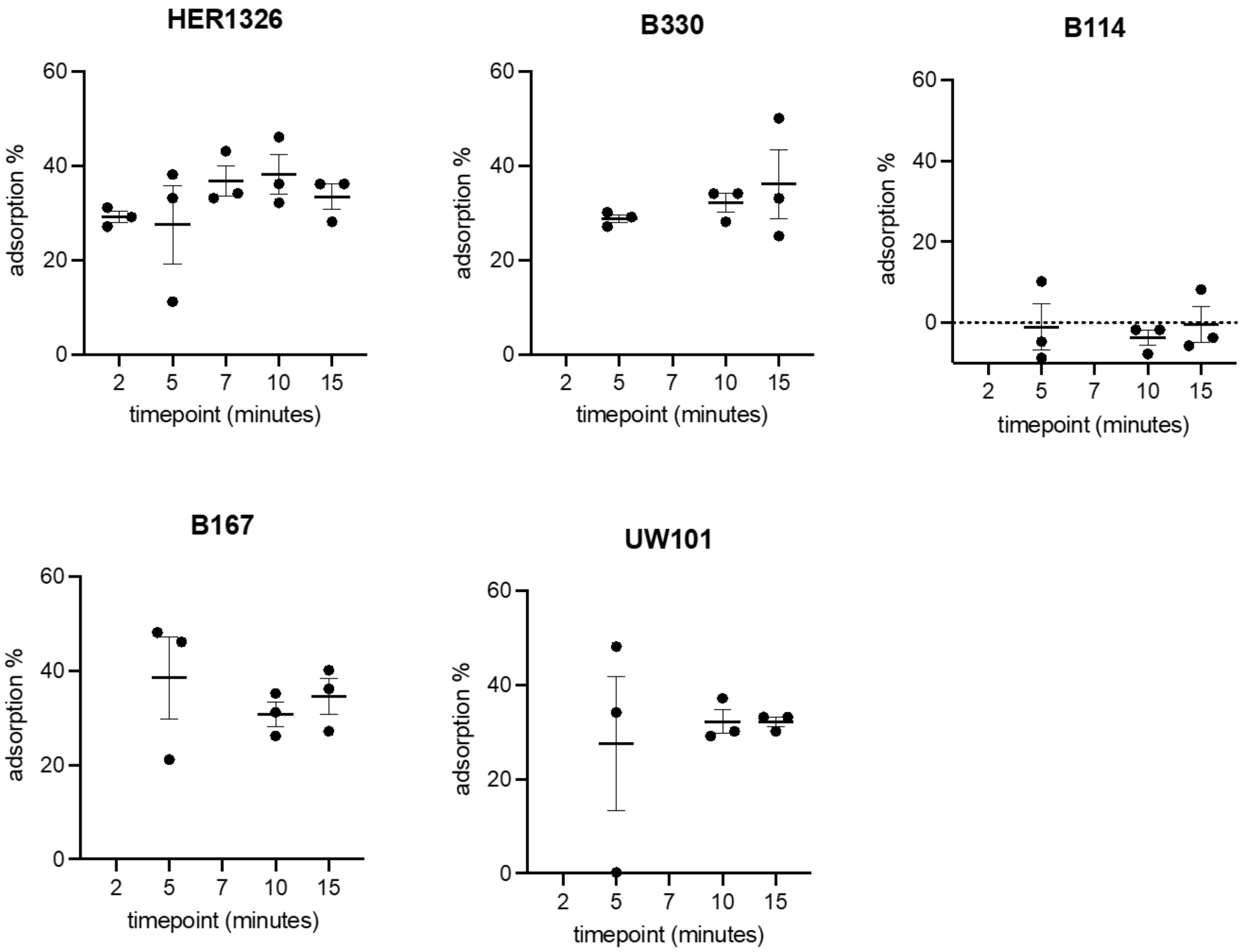

### Extended Data Figure 9

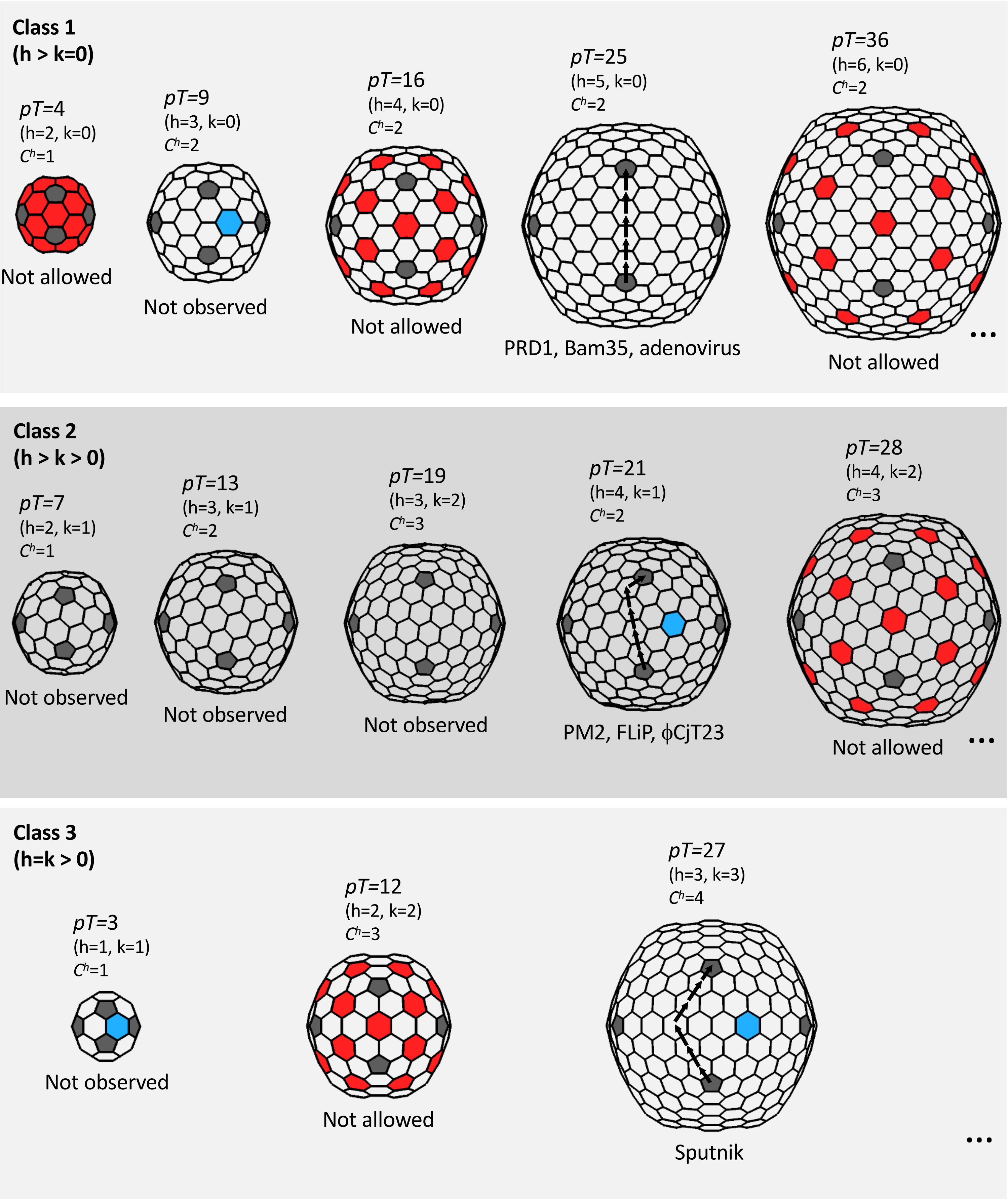
